# Systematic optimization of automated phosphopeptide enrichment for high-sensitivity phosphoproteomics

**DOI:** 10.1101/2023.11.23.568418

**Authors:** Patricia Bortel, Ilaria Piga, Claire Koenig, Christopher Gerner, Ana Martinez del Val, Jesper V. Olsen

## Abstract

Improving coverage, robustness and sensitivity is crucial for routine phosphoproteomics analysis by single-shot liquid chromatography tandem mass spectrometry (LC-MS/MS) runs from minimal peptide inputs. Here, we systematically optimized key experimental parameters for automated on-beads phosphoproteomics sample preparation with focus on low input samples. Assessing the number of identified phosphopeptides, enrichment efficiency, site localization scores and relative enrichment of multiply-phosphorylated peptides pinpointed critical variables influencing the resulting phosphoproteome. Optimizing glycolic acid concentration in the loading buffer, percentage of ammonium hydroxide in the elution buffer, peptide-to-beads ratio, binding time, sample and loading buffer volumes, allowed us to confidently identify >16,000 phosphopeptides in half-an-hour LC-MS/MS on an Orbitrap Exploris 480 using 30 µg of peptides as starting material. Furthermore, we evaluated how sequential enrichment can boost phosphoproteome coverage and showed that pooling fractions into a single LC-MS/MS analysis increased the depth. We also present an alternative phosphopeptide enrichment strategy based on stepwise addition of beads thereby boosting phosphoproteome coverage by 20%. Finally, we applied our optimized strategy to evaluate phosphoproteome depth with the Orbitrap Astral MS using a cell dilution series and were able to identify >32,000 phosphopeptides from 0.5 million HeLa cells in half-an-hour LC-MS/MS using narrow-window data-independent acquisition (nDIA).

**Graphical abstract:** 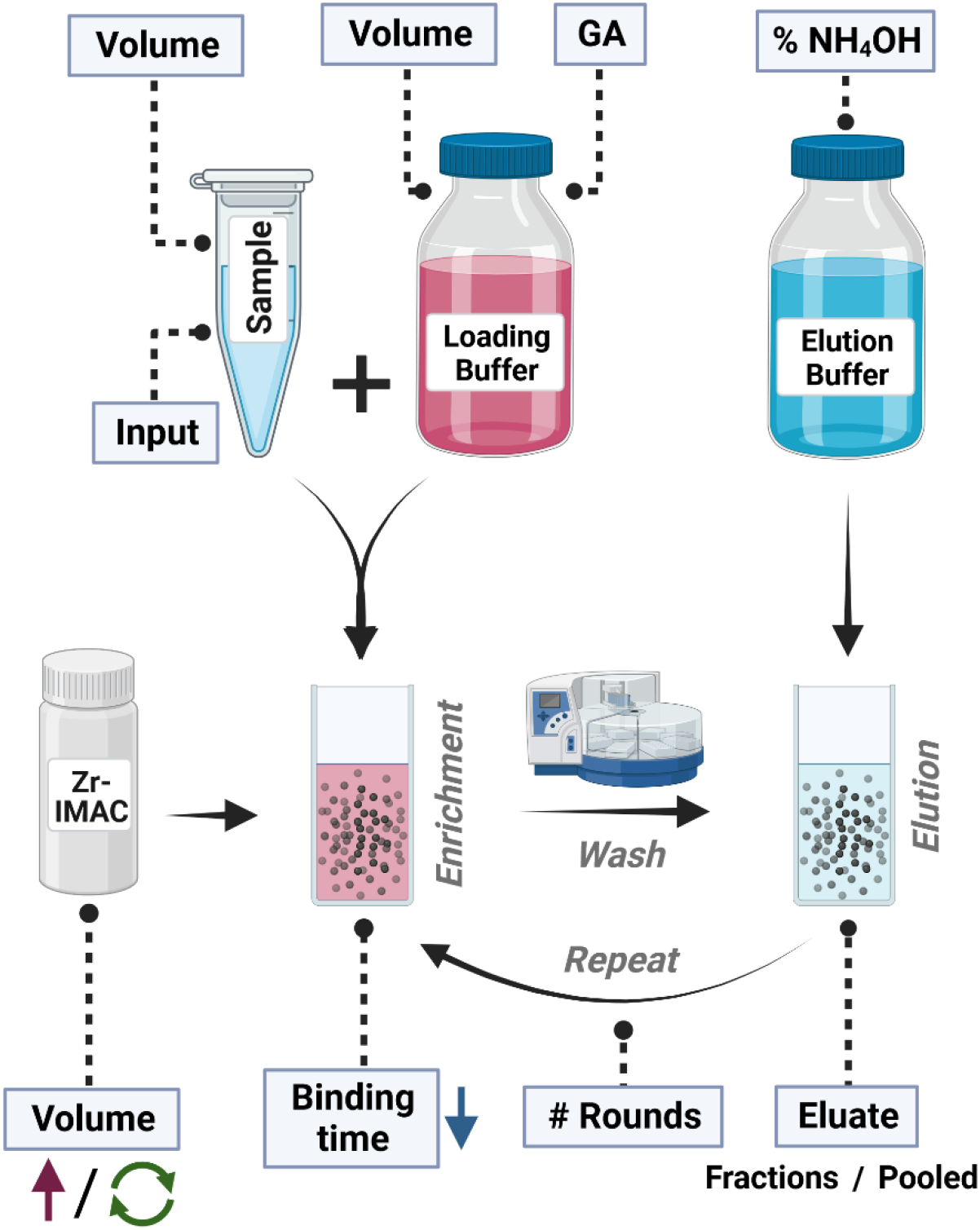

## Introduction

Protein phosphorylation is a highly dynamic post-translational modification (PTM) that plays a critical role in regulating cellular signal transduction pathways. Protein phosphorylation has been the objective of extensive studies by the mass spectrometry (MS)-based proteomics community, with the analysis of thousands of phosphorylation sites across different cellular contexts.^1–5^ However, there is still ongoing research to try to identify the functionality of most of them^6^ since it has been suggested that 75% of the proteome might be phosphorylated^7^. Quantitative mass spectrometry has proved to be the best platform to retrieve large scale information about identification, quantification and localization of phosphorylation sites in complex systems.^5^ The very deep proteomes described so far^8^ or recent development of highly sensitive mass spectrometers^9^ indicate the potential to explore the phosphoproteome without the need for specific enrichment of phosphopeptides prior to MS analysis. However, the capacity to explore the functional phosphoproteome is still impacted by the relatively low abundance of phosphorylated peptides and their sub-stoichiometric nature.^10^ Therefore, to this day, deep phosphoproteomics relies on enrichment of phosphopeptides prior to LC-MS/MS analysis. In this regard, the phosphoproteomics technology has taken a significant leap in recent years with significant improvements in sensitivity, making it possible to perform phosphoproteomics analysis of minute samples as low as single spheroids^11^. Moreover, incorporation of magnetic beads into workflows on automated sample preparation platforms nowadays allows for robust high-throughput sample preparation, making phosphoproteomics applicable to large-scale studies^12–14^ and clinical sample cohorts^15^.

Currently, the most popular phosphopeptide enrichment strategies rely on affinity-based methods either by immobilized metal affinity chromatography (IMAC)^16^ or metal oxide affinity chromatography (MOAC)^17,18^. Both, IMAC and MOAC strategies rely on transition metals (Ti, Zr, Fe) that, either chelated on a substrate (Ti-IMAC or Zr-IMAC) or as metal oxides (TiO2), enable the selective binding of phosphopeptides. The effectiveness of these strategies relies on multiple factors, including the ratio of peptide-to-beads (binding capacity)^19^, the loading buffer composition (binding conditions), the washing buffer composition, and sample-bead binding time, among others^20,21^. For instance, the inclusion of competitive non-phosphopeptide binders, such as glycolic acid (GA) or 2,5-dihydoxybenzoic acid (DHB) in the binding buffer, as well as a high concentration of trifluoroacetic acid (TFA), can minimize the binding of highly-acidic or sialic-acid containing peptides^22–24^, significantly increasing the phosphopeptide enrichment efficiency of IMAC and MOAC strategies. Moreover, different strategies based on the binding of phosphopeptides to the metal matrix have been presented to increase the depth and diversity of the purified phosphopeptide population. For instance, Thingholm et al.^25^ showed that multiply-phosphorylated peptides can be separately purified by sequential elution, based on the higher affinity between the metal matrix and peptides with several phosphate groups. On the other hand, the combination of different bead types has been suggested as a method to enrich complementary sets of phosphopeptides based on their multiplicity and acidity.^26^

Moreover, with the advent of Data Independent Acquisition (DIA) strategies and the implementation of software tools capable of analyzing phosphoproteomics data without the need for spectral libraries^27,28^, the depth obtained from single-shot phosphoproteomics analyses has increased significantly. The combination of short LC gradients with single-shot DIA nowadays provides analysis of deep (phospho)proteomes in a high-throughput manner. In this regard, the improvements achieved by technical sample preparation optimizations have been overshadowed by the increased sensitivity and coverage of detected phosphopeptides by DIA approaches.

In this work, we systematically evaluated how key experimental parameters affect phosphopeptide enrichment with special focus on low input samples, which were subsequently analyzed using DIA. We specifically tested the impact of using different (i) bead-to-peptide ratios (ii) sample-bead-binding times (iii) concentrations of glycolic acid in the loading buffer, (iv) percentages of ammonium hydroxide (NH_4_OH) in the elution buffer, (v) peptide input amounts, (vi) sample volumes and (vii) loading buffer volumes. To identify the optimal phosphopeptide enrichment conditions, the effects of the different parameters were evaluated in terms of phosphopeptide enrichment efficiency, relative purification of multiply-phosphorylated peptides and coverage of well localized phosphosites (class I phosphosites, localization probability >0.75) as a proxy for the quality of the DIA-MS/MS spectra. Then, based on the optimized experimental parameters, we devised different strategies to increase the phosphoproteome depth of the analysis by sequential enrichment strategies, either by repetitive enrichment using the non-bound fraction or by modifying the peptide-to-bead ratio. Finally, we applied our optimized strategy for the phosphoproteomics analysis of a cell dilution series on a state-of-the-art Orbitrap-Astral MS to determine the limits of deep phosphoproteomics analysis with strongly downscaled cell input amounts.

## Methods

### Cell culture and cell lysis

A549 and HeLa cells were cultured in P15 dishes in DMEM (Gibco, Invitrogen) supplemented with 10% fetal bovine serum (FBS, Gibco) and 100 μg/mL streptomycin (Invitrogen) until 90% confluence was reached.

A549 cells were washed twice with phosphate buffered saline (PBS) (Gibco, Life Technologies) and lysed using 600 µL 95°C hot lysis buffer (5% sodium dodecyl sulfate (SDS), 5mM Tris(2-carboxyethyl)phosphine (TCEP), 10mM chloroacetamide (CAA), 100mM Tris pH 8.5). Cells were scraped, collected in a falcon tube and the lysate was incubated at 95°C, 500 rpm for 10 min.

HeLa cells were first detached with 0.05% trypsin-EDTA (Gibco, Invitrogen) and counted *via* a trypan blue viability assay (10 µL of cell suspension was diluted in 1:1 (v/v) ratio with trypan blue stain 0.4% (v/v)) by using an automated cell counter (Corning®). For cell count estimation, the average count of five images acquired with the CytoSMART software was calculated. Cell dilutions with respectively; 1×10^6^, 0.5×10^6^, 0.2×10^6^, 0.1×10^6^, 0.05×10^6^ and 0.01×10^6^ cells were collected in Eppendorf tubes with 4 replicates for each condition. Cell pellets were first washed with PBS and then lysed with 50 µl of 95°C hot SDS 5% and incubated at 95°C, 500 rpm for 10 min. Cell pellets were washed with PBS and, lysed with 50 µL of 95°C hot lysis buffer and incubated at 95°C, 500 rpm for 10 min.

The lysates were cooled to room temperature and sonicated by microtip probe sonication (Vibra-Cell VCX130, Sonics, Newtown, CT). Sonication parameters were set to a total runtime of 2 min with pulses of 1 sec on and 1 sec off at an amplitude of 80%.

### Determination of protein concentration *via* BCA-assay

Protein concentration was determined utilizing the Pierce™ BCA Protein Assay Kit (Thermo Scientific^TM^) according to the manufacturer’s protocol for 96 well plates setup.

For Astral data, mean peptide input amounts were estimated based on 25% recovery after PAC digestion from BCA measurement of protein concentration.

### Automatized Protein Aggregation Capture (PAC) based protein digestion

Protein digestion was performed according to an adapted version of the Protein Aggregation Capture (PAC) based digestion^29^ on a KingFisher^TM^ Flex System (Thermo Scientific^TM^)^30^ with MagReSyn® Hydroxyl beads (ReSyn Biosciences). KingFisher deep-well plates were prepared for washing steps, containing 1 mL of 95% Acetonitrile (ACN) or 70% Ethanol (EtOH). For each sample, 300 µL of digestion solution (50 mM ammonium bicarbonate (ABC)-buffer) containing Lys-C and Trypsin at an enzyme to substrate ratio of 1:500 and 1:250, respectively, were prepared and transferred to KingFisher plates. Samples were mixed with 100 mM Tris-buffer to obtain a total volume of 300 µL, transferred to KingFisher plates and ACN was added to a final volume percentage of 70%. The storage solution from the Hydroxyl beads was replaced with 70% ACN. Finally, beads were added to the samples at a protein to beads ratio of 1:2. Protein aggregation was carried out in two steps of 1 min mixing at medium speed, followed by a 10 min pause each. Sequential washes were performed in 2.5 min at slow speed without releasing the beads from the magnet. Digestion was set to 100 cycles of agitation for 45 seconds and pausing of 6 minutes overnight at 37°C. Protease activity was quenched by acidification with trifluoroacetic acid (TFA) to a final volume percentage of 1%. Processing of HeLa samples was performed similarly, but with some adaptations. The ratio of MagReSyn® hydroxyl beads to protein was 16:1, the time of digestion was 6 hours in 200 ul of 50 mM triethylammonium bicarbonate and the Lys-C and Trypsin to substrate ratio was 1:100 and 1:50 respectively. After digestion, samples were acidified with 50 ul of 10% formic acid, peptides were concentrated in SpeedVac at 45°C until 20 ul and directly processed for phosphopeptide enrichment without Sep-Pak desalting.

### Sep-Pak desalting

Desalting was performed on Sep-Pak C18 cartridges (C18 Classic Cartridge, Waters, Milford, MA). The cartridges were conditioned with 900 µL 100% ACN and 3x 900 µL 0.1% TFA followed by sample loading, and washing 3x with 900 µL 0.1% TFA. Peptides were eluted with 150 µL 40% ACN followed by 150 µL 60% ACN. The acetonitrile was evaporated in a SpeedVac at 45°C and peptide concentration was determined by measuring absorbance at 280 nm on a NanoDrop 2000C spectrophotometer (Thermo Fisher Scientific).

### Phosphopeptide enrichment

Phosphopeptide enrichment was performed on a KingFisher^TM^ Flex System (Thermo Scientific^TM^) using MagReSyn® zirconium-based immobilized metal affinity chromatography (Zr-IMAC HP) beads (ReSyn Biosciences).

#### Standard phosphopeptide enrichment workflow

Samples were mixed with 200 µL loading buffer (80% ACN, 5% TFA, 0.1 M glycolic acid) and transferred to a KingFisher 96 deep-well plate. Additional KingFisher plates were prepared containing 500 µL of loading buffer, 500 µL of washing buffer 2 (80% ACN, 1% TFA) or 500 µL of washing buffer 3 (10% ACN, 0.2% TFA) each. For each sample, 5 µL of beads (20 mg/mL) were suspended in 500 µL 100% ACN previously added to the KingFisher plates. For elution, 200 µL of elution buffer (1% NH_4_OH) were prepared and transferred to KingFisher plates. Beads were washed in loading buffer for 5 min at 1000 rpm, incubated with the samples for 20 min with mixing at medium speed and subsequently washed in loading buffer, washing buffer 2 and washing buffer 3 for 2 min with mixing at fast speed. Phosphopeptides were eluted from the beads by mixing with elution buffer for 10 min at fast speed.

When evaluating the effect of different experimental parameters, the standard phosphopeptide enrichment workflow was altered as indicated in the experimental design table 1. For evaluating the influence of varying sample volume while keeping the peptide input the same, samples were diluted with 1% TFA to the final desired concentrations.

**Table 1.** Experimental design for the optimization of phosphopeptide enrichment parameters. Evaluated parameters included peptide input (Peptide µg), sample volume (Sample V), loading buffer volume (LB V), concentration of glycolic acid in the loading buffer (GA in LB), Zr-IMAC HP bead volume (Bead V), beads binding time (Bead binding) and percentage of ammonium hydroxide in the elution buffer (NH_4_OH in EB).

| Parameter optimization | Peptide input [μg] | Sample V [μl] | LB V [μl] | GA in LB [mol/l] | Beads V [μl] | Bead binding [min] | NH<sub>4</sub>OH in EB [% (v/v)] |
| --- | --- | --- | --- | --- | --- | --- | --- |
| Standard | 30 | 3.29 | 200 | 0.1 | 5 | 20 | 1 |
| Peptide μg | 5-30 | Standard | Standard | Standard | Standard | Standard | Standard |
| Sample V | Standard | 7.5-120 | Standard | Standard | Standard | Standard | Standard |
| LB V | Standard | 15 | 100-400 | Standard | Standard | Standard | Standard |
| GA in LB | Standard | Standard | Standard | 0-2 | Standard | Standard | Standard |
| Bead V | Standard | Standard | Standard | Standard | 1-20 | Standard | Standard |
| Bead binding | Standard | Standard | Standard | Standard | Standard | 5-30 | Standard |
| NH<sub>4</sub>OH in EB | Standard | Standard | Standard | Standard | Standard | Standard | 0.1-2 |

#### Standard sequential/ Looped enrichment workflow

Standard sequential enrichment was performed for 2-3 enrichment rounds by re-starting the phosphopeptide enrichment workflow on the KingFisher^TM^ Flex System while keeping samples, beads and washing buffers the same. For obtaining “pooled” samples from the sequential enrichment, all rounds were carried out re-using the same elution buffer. In order to retrieve the phosphopeptides captured in each round as separate fractions, the elution buffer was removed after each round and replaced by new elution buffer.

#### Sequential phosphopeptide enrichment workflow adaptations

Parameters of the sequential phosphopeptide enrichment workflow were altered as indicated in the experimental design table 2. For sequential enrichment with increasing molarity of glycolic acid in the loading buffer, 64 µL loading buffer containing 8 M glycolic acid (GA) were added to the sample in 200 µL standard (0.1 M GA) loading buffer after the first round of enrichment to adjust to 2 M glycolic acid. Sequential enrichment with additional Zr-IMAC HP beads per round was performed by addition of 1 µL or 2 µL Zr-IMAC HP beads after the first and second round, respectively, to the beads plate containing 1 µL starting amount of beads. Sample-bead incubation times of 1 min and 5 min were applied for testing sequential enrichment in combination with short incubation times. For testing the effect of fresh beads in a sequential approach, Zr-IMAC HP beads were removed after the first round of enrichment, and replaced by the same amount of fresh beads for a second enrichment step.

**Table 2.**
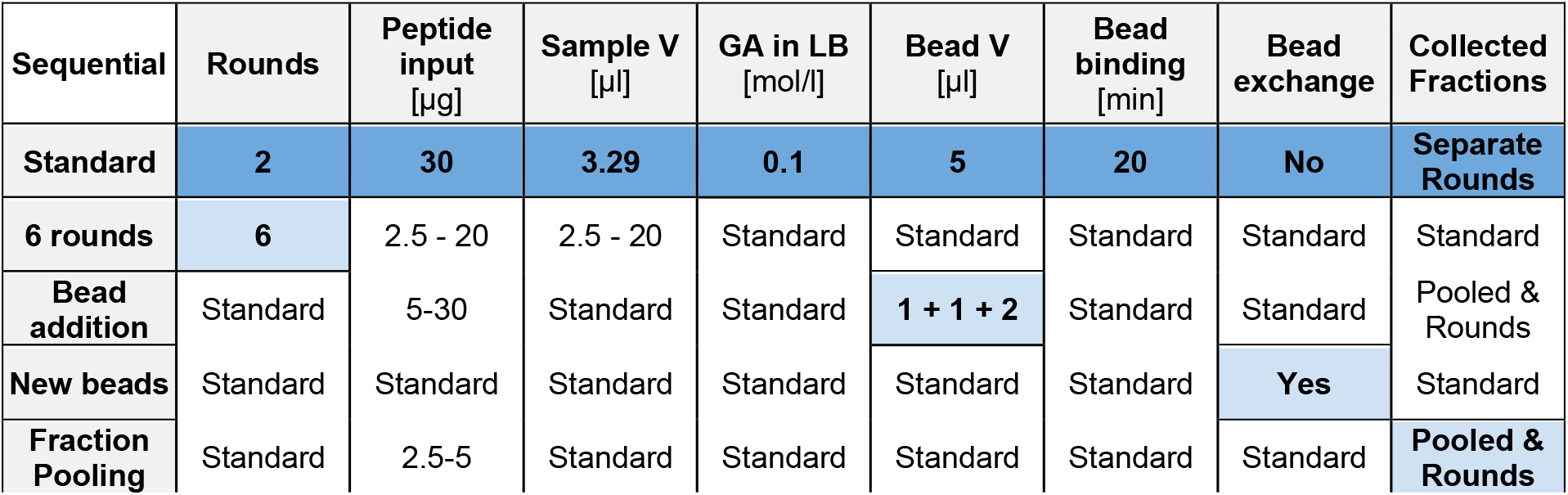
Experimental design for the optimization of sequential phosphopeptide enrichment. The sequential enrichment workflow design was evaluated in terms of reusing or exchanging the beads after the first enrichment round (New beads), total number of enrichment rounds (6 rounds), addition of beads with increasing enrichment round (Bead addition), and sample pooling approaches for LC-MS/MS analysis (Fractions).

| Sequential | Rounds | Peptide input<br>[μg] | Sample V<br>[μl] | GA in LB<br>[mol/l] | Bead V<br>[μl] | Bead binding<br>[min] | Bead exchange | Collected Fractions |
| --- | --- | --- | --- | --- | --- | --- | --- | --- |
| Standard | 2 | 30 | 3.29 | 0.1 | 5 | 20 | No | Separate Rounds |
| 6 rounds | 6 | 2.5 - 20 | 2.5 - 20 | Standard | Standard | Standard | Standard | Standard |
| Bead addition | Standard | 5-30 | Standard | Standard | 1 + 1 + 2 | Standard | Standard | Pooled & Rounds |
| New beads | Standard | Standard | Standard | Standard | Standard | Standard | Yes | Standard |
| Fraction Pooling | Standard | 2.5-5 | Standard | Standard | Standard | Standard | Standard | Pooled & Rounds |

### Sample preparation for LC-MS/MS analysis

Eluates containing phosphopeptides were acidified with 40 µL 10% TFA to achieve a pH <2. Acidified eluates were transferred to MultiScreen®_HTS_-HV 96-well filtration plates (0.45 μm, clear, non-sterile, Millipore), stacked on 96-well plates and centrifuged for 1 min at 500 ×g to remove in-suspension particles.

Evotip Pure^TM^ (Evosep) were washed by adding 20 µL 100% ACN and centrifuging for 1 min at 800 ×g. Tips were pre-conditioned by addition of 20 µL 0.1% formic acid (FA) while soaking the tips in 100% isopropanol and centrifuged for 1 min at 800 ×g. Filtered samples were added to the tips and loaded by centrifugation for 2 min at 500 ×g. Evotip preparation was completed by adding 20 µL of 0.1% FA, centrifuging for 1 min at 800 ×g, adding 200 µL of 0.1% FA and centrifuging for 10 s at 800 ×g.

### LC-MS/MS analysis

Samples were analyzed using an IonOpticks Aurora™ column (15cm-75μm-C_18_ 1.6μm) interfaced with the Orbitrap Exploris™ 480 Mass Spectrometer (Thermo Scientific™) or the Orbitrap Astral Mass Spectrometer (Thermo Scientific™) using a Nanospray Flex™ Ion Source with an integrated column oven (PRSO-V2, Sonation, Biberach, Germany) to maintain the temperature at 50 °C. In all samples, spray voltage was set to 1.8 kV, funnel RF level at 40, and heated capillary temperature at 275 °C. Samples were separated on an Evosep One LC system using the pre-programmed gradient for 40 samples per day (SPD).

For phosphoproteome analysis of A549 samples using DIA on the Orbitrap Exploris™ 480 Mass Spectrometer, full MS resolutions were set to 120,000 at m/z 200 and the full MS AGC target was 300% with an IT of 45 ms. The AGC target value for fragment spectra was set to 1000%. 49 windows of 13.7 m/z scanning from 472 to 1143 m/z were employed with an overlap of 1 Da. MS2 resolution was set to 15,000, IT to 22 ms and normalized collision energy (NCE) to 27%.

For phosphoproteome analysis of HeLa samples using DIA on the Orbitrap Astral MS. The MS was operated at a full MS resolution of 180,000 with a full scan range of 480 − 1080 *m/z*. The full scan AGC target was set to 500%. Fragment ion scans were recorded at a fixed resolution of 80,000 and with a maximum injection time of 6 ms. 150 windows of 4 m/z scanning from 380-980 m/z were used. The isolated ions were fragmented using HCD with 27% NCE.

### Data analysis

LC-MS/MS runs were searched using Spectronaut (version 17.1 for Orbitrap Exploris 480 data and version 18 for Orbitrap Astral data) employing a directDIA^TM^ search strategy against the homo sapiens proteome UniProt Database (2022 version, 20,958 entries) supplemented with a database of common contaminants (246 entries). Carbamidomethylation of cysteine was set as fixed modification, whereas oxidation of methionine, N-terminal protein acetylation and phosphorylation of serine, threonine and tyrosine were set as variable modifications. The maximum number of variable modifications per peptide was set to 5, cross-run normalization was turned off and method evaluation was turned on. PTM localization was turned on, but the localization probability threshold was set to 0.

Searches of data from sequential enrichment sets were performed by searching the enrichment rounds and/or fractions separately, while keeping replicates of the same peptide input amount in the same analysis.

Precursor level pivot tables were exported from Spectronaut for phosphopeptide reporting analysis. Tables were filtered to contain unique modified sequences (i.e. phosphopeptide isomers – same stripped sequence but different phosphorylation site – are kept as separated entities) and unique modified sequences containing phosphorylation sites were further filtered to preserve only those with a localization score >0.75 in at least one replicate.

Data was exported in long-format and imported into Perseus (v1.6.5.0) where it was collapsed into phospho-sites or phosphopeptides using the “peptide-collapse” plugin (v1.4.2) described in Bekker-Jensen et al.^31^. For collapse into phosphosites, the option “Target PTM site-level” was used. By default, the localization cutoff was kept at 0.75. When evaluating the localization probability dsitribution, the localization cutoff was set to 0. Importantly, phopshosites reported by “peptide-collapse” plugin must have been identified and/or localized in at least two experimental replicates. Collapse into phosphopeptides using Perseus plugin was employed for quantification purposes. For phosphopeptide collapse, the option “ModSpec peptide-level” was used and localization cutoff was kept at 0.75. Phosphopeptide collapse in Perseus will grouped together different phospho-isomers.

All remaining processing steps were performed either in Perseus (v1.6.5.0) or R (v3.6.2 or higher) implementing the packages ComplexHeatmap^32^, sitools^33^, eulerr^34^, stringr^35^, ggplot2^36^, dplyr^37^, tidyverse^38^ and limma^39^. Calculation of isoelectric point (pI) values was performed using the package pIR^40^, considering N-terminal acetylation and phosphorylation of the peptides.

## Results and discussion

### Phosphopeptide binding conditions affect the population of purified phosphopeptides

To establish an optimized automated phosphopeptide enrichment procedure for high-sensitivity samples^12^, we started by evaluating the main experimental parameters that can affect the phosphopeptide enrichment. This was done by introducing modifications to our default automated protocol for sensitive phosphoproteomics, which relies on the use of magnetic Zr-IMAC HP beads in the KingFisher^TM^ System. The resulting phosphopeptide mixtures were subsequently analyzed by DIA-MS in an Orbitrap Exploris 480 MS coupled to an Evosep One LC system taking advantage of the higher sensitivity of the Whisper gradients. All enrichments were performed from a starting amount of 30 µg of purified peptides from a whole cell tryptic digest of the A549 lung cancer cell line, as it represents the optimal peptide input amount for phosphopeptide enrichment and subsequent analysis with our LC-MS/MS setup using flow Whisper gradients. By using this amount, we ensured to have an adequate reference to assess the effect of the changes in the experimental workflow. The phosphopeptide enrichment protocol consists of three main steps: (i) the binding of phosphopeptides to the beads, (ii) the washing of the beads to remove non-specific interactions and (iii) the elution of phosphopeptides from the beads (**Figure 1A**). We focused on the first part of the protocol, the binding of phosphopeptides to the beads, in which we evaluated the following parameters: (i) the beads to peptide ratio, (ii) the proportion of glycolic acid in the loading buffer, (iii) the binding time and (iv) the sample to loading buffer volume ratio. In the last step of the protocol, we evaluated the effect of modifying the concentration of ammonium hydroxide in the elution buffer (**Figure 1B and Table 1**).

**Figure 1.**
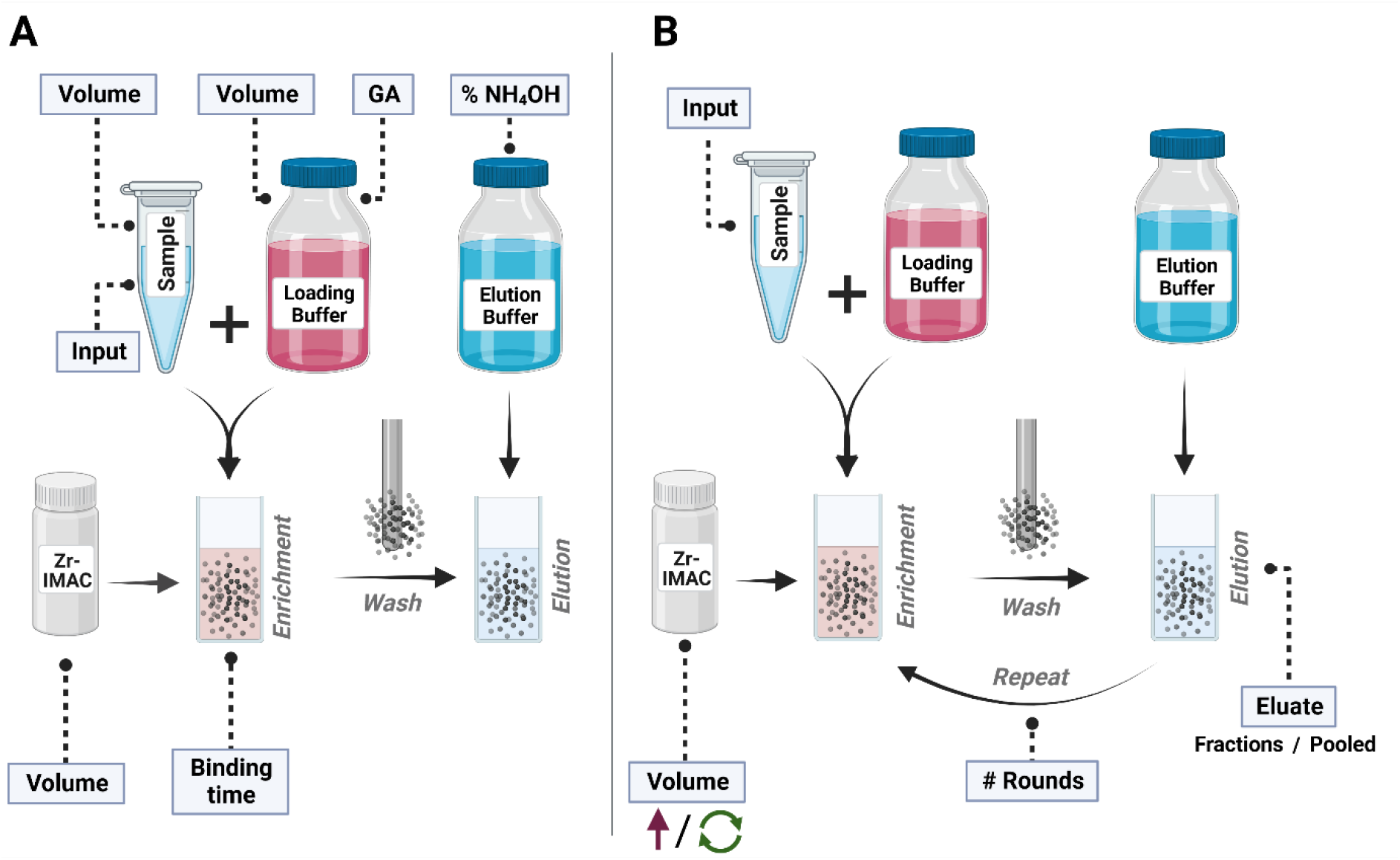
Experimental design. **(A)** Schematic overview of the phosphopeptide enrichment workflow and the evaluated experimental parameters. Evaluated parameters included the peptide input, the sample volume, the loading buffer volume, the proportion of glycolic acid in the loading buffer, the percentage of ammonium hydroxide in the elution buffer, the Zr-IMAC HP bead volume and the sample-beads binding time. **(B)** Schematic overview of the sequential phosphopeptide enrichment workflow and the evaluated experimental parameters. Evaluated parameters included the peptide input amount of the sample, the number of sequential enrichment rounds and the way of retrieving the eluate by either reusing the elution buffer and obtaining a “Pooled” eluate or exchanging the elution buffer after each round and obtaining each enrichment round as single fraction for LC-MS/MS analysis. Modified sequential enrichment approaches included exchanging, reusing or increasing the Zr-IMAC HP bead volume.

The proteomics community has extensively evaluated the beads-to-peptide ratio as this plays an important role for the phosphopeptide enrichment efficiency. A high beads-to-peptide ratio can lead to increased binding of non-phosphorylated peptides due to nonspecific interactions with the bead surface, thus reducing the selectivity of the enrichment. In contrast, it has also been described that a too low beads-to-peptide ratio can result in a higher fraction of multiply-phosphorylated peptides identified.^19^ In this work, we assessed the effect of using different bead amounts. This is particularly relevant when performing enrichment with low peptide amounts, since it is difficult to proportionally scale down the volume of beads, due to lack of reproducibility while pipetting low volumes of beads. Therefore, starting from 30 µg of peptides, we evaluated what the best compromise between bead volume and good phosphopeptide recovery would be by testing 1, 2, 5, 10 and 20 µL of beads corresponding to a beads-to-peptide ratio of 0.7, 1.3, 3.3, 6.7 and 13.3 (**Figure 2A**). We observed that the best outcome was obtained using a volume of 5 µL (beads-to-peptide ratio of 3.3), which resulted in 16,193 phosphopeptides. Importantly, throughout this work we only report a phosphopeptide as valid for those where the phosphorylation was localized to an amino acid with a score > 0.75. Increasing the bead volume to 20 µL slightly decreased the overall number of phosphopeptides to 15,026 as well as the relative enrichment efficiency based on counts (i.e., number of phosphopeptides versus total number of peptides) (from 79 % with 5 µL to 62 % with 20 µL). On the other hand, reducing the bead volume to 1 µL resulted in higher enrichment efficiency (84 %) (**Figure 2E**) and trend towards of more acidic phosphopeptides (**Figure 2H**), although lower overall phosphoproteome coverage (12,962 phosphopeptides) (**Figure 2A-F**). Our titration experiment also confirmed that the selectivity of the enrichment in regard to the purification of mono- or multiply-phosphorylated peptides is affected by the available binding surface, as the absolute number of multiply-phosphorylated peptides increased with decreasing bead amount (**Figure 2G-I**).

**Figure 2.**
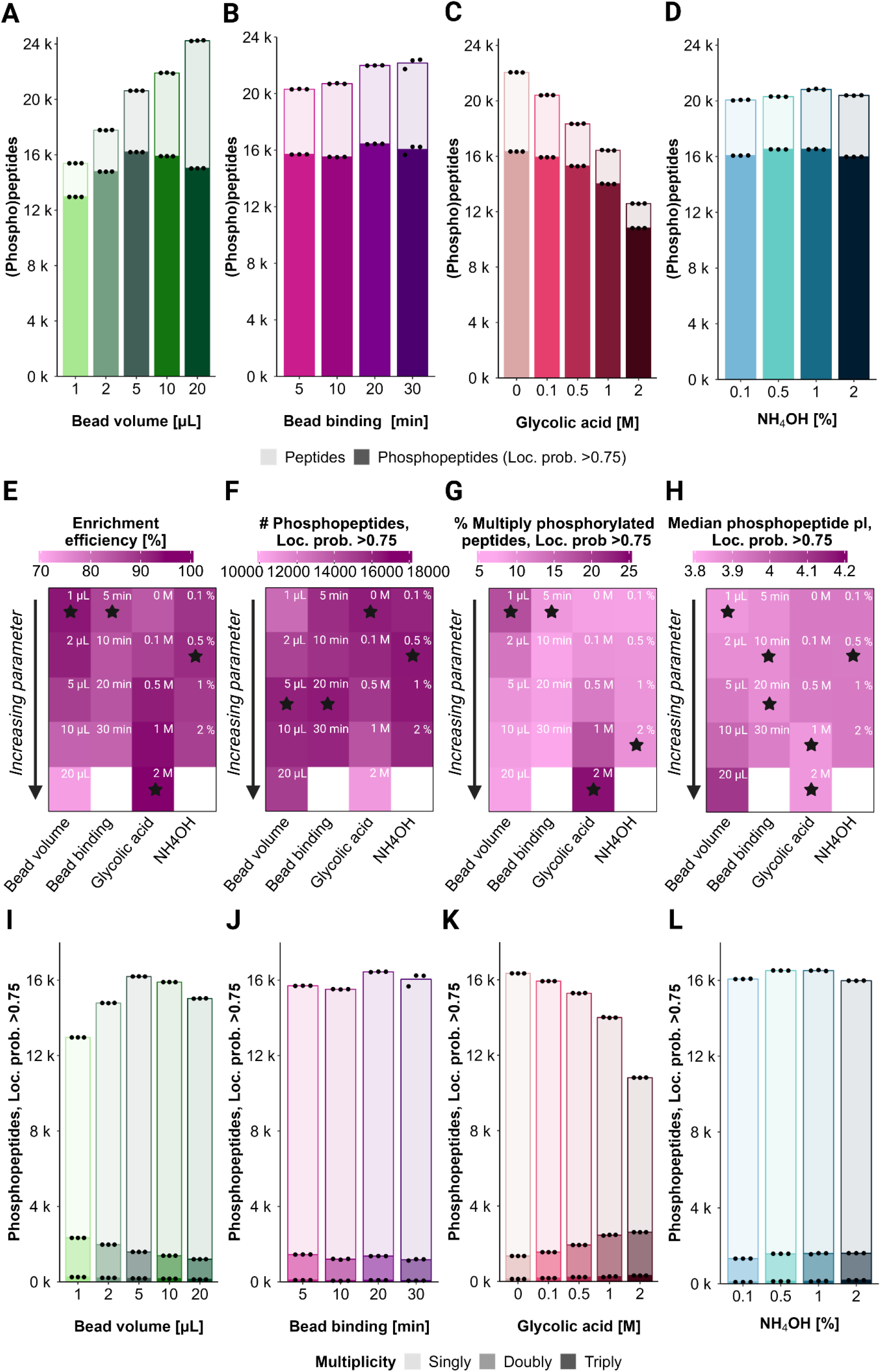
Evaluation of experimental parameters during phosphopeptide enrichment. **(A-D)** Barplots show the mean numbers of peptides (light color) and phosphopeptides with loc. prob. >0.75 (dark color) identified across three experimental replicates using different **(A)** molarities of glycolic in the loading buffer, **(B)** percentage of ammonium hydroxide in the elution buffer, **(C)** Zr-IMAC HP bead volumes or **(D)** sample-bead binding times. Each dot represents one experimental replicate. **(E-H)** Heatmaps show the influence of increasing molarity of glycolic acid in the loading buffer, percentage of ammonium hydroxide in the elution buffer, Zr-IMAC HP bead volume or sample-bead incubation time on **(E)** selectivity of the enrichment in terms of identified phosphorylated and non-phosphorylated peptides **(F)** number of identified phosphopeptides with loc. prob. >0.75 **(G)** percentage of multiply-phosphorylated peptides with loc. prob. >0.75 in context of total identified phosphorylated peptides with loc. prob. >0.75 **(H)** median pI of phosphopeptides with loc. prob. >0.75. Stars refer to the highest value within the respective parameter column for **E-G** and to the lowest value for **H**. If not otherwise indicated, all values represent the mean of three experimental replicates. **(I-L)** Barplots show the mean numbers of singly (light color), doubly (medium color) and triply (dark color) phosphorylated peptides with loc. prob. >0.75 identified across three experimental replicates using different **(I)** molarities of glycolic in the loading buffer, **(J)** percentage of ammonium hydroxide in the elution buffer, **(K)** Zr-IMAC HP bead volumes or **(L)** sample-bead binding times. Each dot indicates one experimental replicate.

We next evaluated the effect of changing the beads-to-peptide binding time to explore whether shortening the binding time would lead to lower phosphoproteome depth and/or bias the recovery towards multiply-phosphorylated peptides. Shortening the binding time from 20 to 5 minutes had a slight impact on the phosphoproteome depth achieved, with 15,700 phosphopeptides quantified after 5 minutes binding compared to 16,440 after 20 min binding (**Figure 2B-F**). However, it seems that there was a slight improvement in enrichment efficiency (from 75 % in 20 minutes to 77 % in 5 minutes), likely due to more unspecific binding to the beads with longer incubation time (**Figure 2E**). Moreover, only a marginal increase in the percentage of multiply-phosphorylated peptides was observed by shortening the binding time (**Figure 2G-J**).

The use of non-phosphopeptide excluders during binding to prevent binding of non-phosphopeptides with high affinity towards IMAC-metal conjugates was evaluated next. In our standard protocol, we originally used 0.1 M of glycolic acid (GA) as a competitive binder in the loading buffer, as recommended by the beads’ manufacturer. With this GA concentration, we obtained an enrichment efficiency of 78 % based on peptide counts or 94% based on MS signal abundance. We observed that increasing the GA concentration up to 2M improved the overall enrichment efficiency (86 % based on peptide counts) and slightly lowered the median phosphopeptide pI (**Figure 2H**), but at the cost of reduced phosphoproteome depth with 10,811 phosphopeptides quantified against 15,929 in our standard protocol (**Figure 2C-E-F**). In contrast, removing the GA greatly decreased the enrichment efficiency (to 74% based on peptide counts), while preserving the phosphoproteome coverage (16,342 phosphopeptides with 0M GA) obtained with 0.1 M of GA (**Figure 2C-E-F**). Interestingly, the high concentrations of GA (1M and 2M) biased the enrichment towards multiply-phosphorylated peptide species, providing up to 16% more multiply-phosphorylated peptides than 0.1M of GA (**Figure 2G-K**). This demonstrated that in high concentrations, GA does not only compete with acidic amino acids in its function as competitive binder, but also with phosphopeptides. Hence, the most competitive phosphopeptides (multiply phosphorylated species) would bind preferentially. Overall, we could confirm that multiply-phosphorylated peptides have a higher affinity towards Zr-IMAC HP beads, explaining why they are preferably recovered with more competitive binding conditions such as 2M GA or shorter binding time.

Next, we questioned whether the lower multiply-phosphorylated peptide recovery in standard enrichment conditions (0.1 M GA, 20 min binding time) could be due to the stronger binding of multiply-phosphorylated peptides to the beads, preventing them from proper elution when present in a large pool of singly-phosphorylated peptides. We evaluated different concentrations of ammonium hydroxide (NH_4_OH) for phosphopeptide elution from the beads, ranging from 0.1 to 2% (v/v). Although subtle, we observed that the lowest NH_4_OH concentration tested (0.1%) resulted in lower multiply-phosphorylated peptide recovery with 8% multiply-phosphorylated phosphorylated peptides compared to 10% multiply-phosphorylated peptides with 2% NH_4_OH (**Figure 2D-G-L**). Enrichment efficiency was highest with 0.5% NH_4_OH (81%) and slightly decreased with higher NH_4_OH concentrations (79% with 1% NH4OH and 78% with 2% NH4OH) (**Figure 2D-E**). Altogether, we conclude that percentage of NH_4_OH in the elution buffer doesn’t significantly impact the elution of bound phosphopeptides from the beads.

To test the influence of the sample input amount in the low peptide range, we performed phosphopeptide enrichment using 30, 10 and 5 µg of peptides. As expected, phosphopeptide recovery is highly dependent on the sample input with 6,888 phosphopeptides quantified from 5 µg compared to 12,480 phosphopeptides quantified using 30 µg of peptide amount (**Figure 3A**). Also, the site localization scores scaled with sample amount (**Figure 3B**), reflecting that the capacity of the search engine to localize phosphorylation sites is dependent on the signal and therefore the quality of the MS2 spectra. The percentage of multiply-phosphorylated peptides barely increased with increasing peptide amount (5% for 5 µg, 6% for 15 µg and 7% for 30 µg) (**Figure 3C**). Finally, phosphopeptides enriched from different peptide input amounts highly overlapped, with more abundant phosphopeptides being preferentially enriched from all input amounts, and the phosphoproteome depth increased as expected with higher input amounts, where lower abundant phosphopeptides were detectable (**Figure 3D, Figure 3E**).

**Figure 3.**
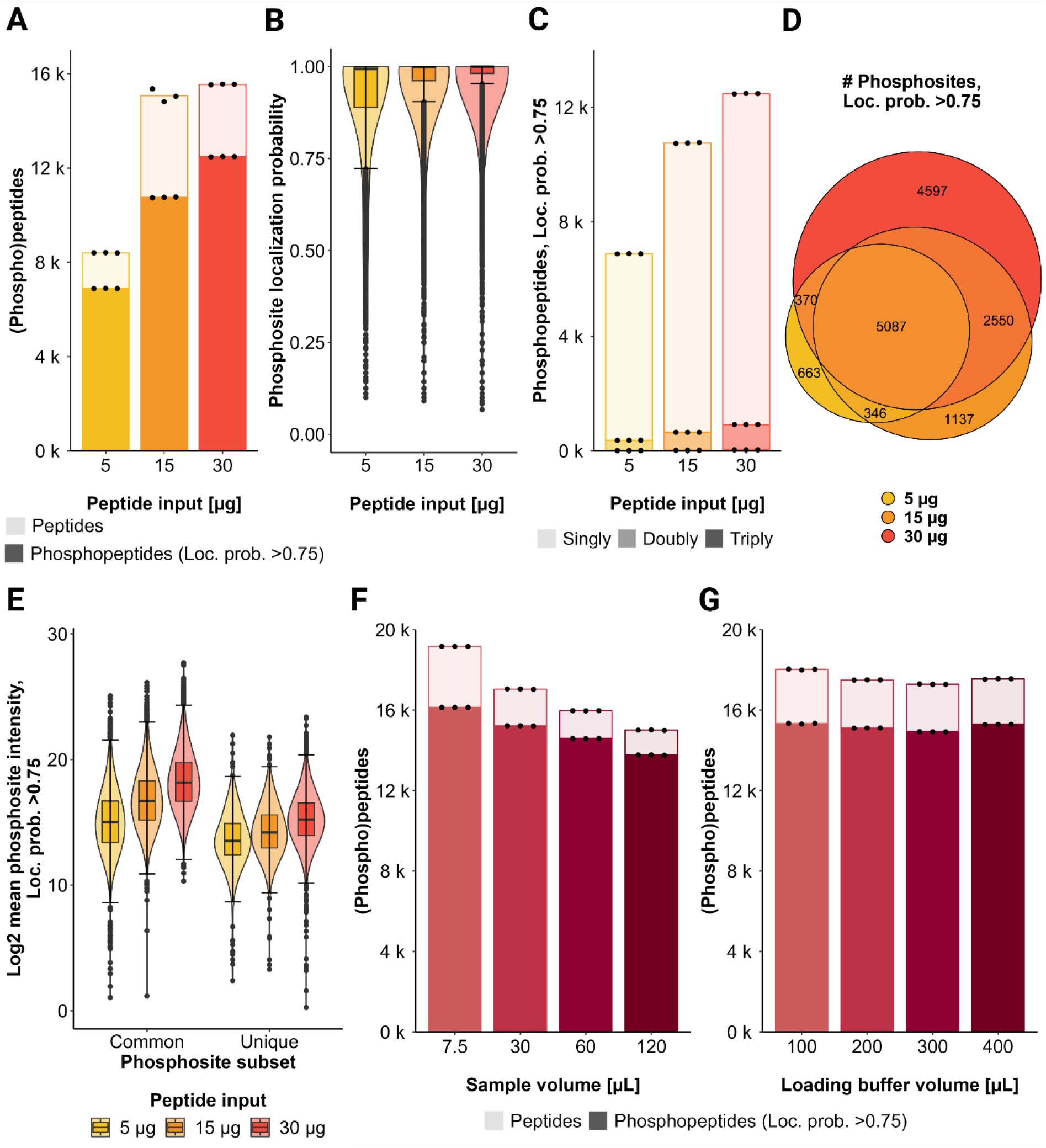
Evaluation of peptide input and sample / loading buffer volume effects. **(A)** Barplots show the mean number of peptides (light color) and phosphopeptides with loc. prob. >0.75 (dark color) identified across three experimental replicates using different peptide input amounts. Each dot represents one experimental replicate. **(B)** Violin plots show the range and distribution of the localization probability of phosphosites identified using different peptide input amounts. **(C)** Barplots show the mean numbers of singly (light color), doubly (medium color) and triply (dark color) phosphorylated peptides with loc. prob. >0.75 identified across three experimental replicates using different peptide input amounts. Each dot represents one experimental replicate. **(D)** The venn diagram shows uniquely and commonly identified phosphosites with loc. prob. >0.75 among different peptide input amounts. **(E)** Violin plots show the log2 mean intensities of uniquely and commonly identified phosphosites with loc. prob. >0.75 among different peptide input amounts. **(F)** Barplots show the mean numbers of peptides (light color) and phosphopeptides with loc. prob. >0.75 (dark color) identified across three experimental replicates using the same peptide input amount (30 µg) diluted in different sample volumes, mixed with the same volume of loading buffer (200 µL). Each dot represents one experimental replicate. (**G**) Barplots show the mean numbers of peptides (light color) and phosphopeptides with loc. prob. >0.75 (dark color) identified across three experimental replicates using the same peptide input amount (30 µg) diluted in the same sample volume (15 µL), mixed with different volumes of loading buffer. Each dot represents one experimental replicate.

Finally, we evaluated the sample volume-to loading buffer (LB) volume ratio. First, we diluted the sample while keeping the LB volume constant (**Table 1 and Figure 3F**), and second, we kept the sample highly concentrated in a constant volume of 15 µL while using increasing LB volumes (**Table 1 and Figure 3G**). Increasing the ratio sample volume-to-LB volume had a negative impact on the phosphoproteome depth (16,132 phosphopeptides with 7.5 µL sample volume vs. 13,770 phosphopeptides with 120 µL sample volume), potentially due to the dilution of the LB by addition of higher sample volumes during binding. However, the dilution of LB in increasing volumes of sample had an impact on the enrichment by reducing the binding of non-phosphorylated species (2,691 non-phosphorylated peptides with 7.5 µL sample volume vs. 2,258 non-phosphorylated peptides with 120 µL sample volume) (**Figure 3F**). On the other hand, we observed that the volume of the loading buffer did not have an impact on the phosphoproteome depth or enrichment efficiency, as long as the sample was kept to a minimal volume (< 30 µL) (**Figure 3G**).

Altogether, in Zr-IMAC HP based phosphopeptide enrichment, we observed that the beads-to-peptide ratio, the concentration of GA in the loading buffer, the peptide input itself as well as the sample concentration can have significant impact on the resulting phosphoproteomes. On the contrary, the binding time and percentage of NH_4_OH in the elution buffer did not seem to have such a significant influence on the phosphopeptide enrichment.

Our evaluation showed that for highly sensitive phosphoproteomics, 5 µL of Zr-IMAC HP beads, 20 min binding time, 0.1 M GA in the loading buffer and 0.5% of NH_4_OH in the elution buffer should be employed to obtain the best phosphopeptide enrichment. However, when multiply phosphorylated peptides are of interest, highly competitive binding conditions such as using 1 µL of Zr-IMAC HP beads, 5min binding time, 2M GA in the loading buffer and a high percentage of NH_4_OH in the elution buffer (2%) could be the best choice.

### Sequential enrichment of the phosphoproteome as a strategy to increase the depth of the analysis

Our data so far showed that changing experimental parameters during phosphopeptide enrichment can have a significant impact on the population of enriched phosphopeptides. Exploring these differences has been suggested before as a potential way to enhance the performance of phosphopeptide enrichment strategies by performing sequential enrichment. The most straightforward way to perform sequential enrichment is to iterate the enrichment by using the flow-through from the previous enrichment. This strategy has been utilized before to increase the depth of the phosphoproteome^13,21,41^. Therefore, we wanted to evaluate how many sequential rounds of enrichment are needed to efficiently deplete a sample for phosphopeptides, and whether sequential enrichment is as efficient with high peptide input amounts as it is with low peptide input amounts.

First, we evaluated whether the beads employed in one round of phosphopeptide enrichment could be reused for a second enrichment. Our data reflected the potential of reusing the beads for sequential enrichment. Interestingly, reusing the beads from the 1^st^ enrichment in a 2^nd^ one resulted in a higher phosphopeptide recovery in the second enrichment round (10,928 phosphopeptides with new beads compared to 12,812 phosphopeptides with reused beads) and a better enrichment efficiency, when compared to using new beads for the second enrichment (52 % with new beads compared to 60 % with reused beads) (**Supplementary Figure S1**).

Next, we tested more extensive sequential enrichment (up to six rounds) from samples spanning 20 to 2.5 µg of peptide input. Whilst the enrichment efficiency (based on phosphopeptide intensities, measured as the percentage of the overall measured MS signal intensity from phosphopeptides alone) was above 90% in the first enrichment round for all amounts tested, it abruptly decreased with lower peptide input amounts in subsequent enrichment rounds down to 86% for 20 µg and 58% for 2.5 µg in round six (**Figure 4A**). Similarly, the phosphoproteome depth (phosphopeptide count relative to round 1) decreased with each sequential enrichment, which was more evident for lower input amounts (**Figure 4A-C-D**). The number of additional unique phosphopeptides did not significantly increase after the third enrichment round (**Figure 4F and Supplementary Fig S2A**), even though up to 2,105 phosphopeptides were still enriched in the sixth enrichment of 20 µg peptide input amount (**Figure 4A-C**). In line with this, the enrichment efficiency based on MS signal intensity, decreased with each subsequent enrichment (**Figure 4A and Supplementary Figure S2A-B**). The population of phosphopeptides enriched in each subsequent enrichment, especially from third and forward, was mainly driven by abundance, since the most abundant peptides in the first enrichment round continued being enriched subsequently (**Supplementary Figure S2A**). Unlike the phosphopeptides, the non-phosphorylated peptides eluted differently across the sequential enrichments (**Supplementary Figure S2A**). At least three different trends were observed in the non-phosphorylated peptides elution: (group 1) peptides eluting mainly in the first fraction (enrichment round), (group 2) peptides eluting mainly in the second fraction and (group 3) peptides with a consistent elution between fractions (**Supplementary Figure S2A**). We decided to explore the nature of these peptides further to understand the mechanisms behind the non-phosphopeptide binding with Zr-IMAC HP beads. When compared to a comprehensive proteome of the same cell line (A549), it was evident that the non-phosphorylated peptides that bound to the beads belonged to the most abundant pool of peptides in the proteome (**Supplementary Figure S2C**). In particular, the non-phosphorylated peptides that showed a consistent elution across the first three fractions (group 3) were more abundant in the proteome than the rest, showing that their binding is most likely mainly driven by abundance (**Supplementary Figure S2C**). We also estimated the isoelectric point of these peptides and found them to be generally acidic (pI ∼5), although less than the phosphopeptides (pI ∼3) (**Supplementary Figure S2D**). Altogether, this data shows that, even though the selectivity will strongly favor the binding of phosphopeptides, the high abundance of non-phosphorylated peptides can lead to unspecific binding during sequential enrichment.

**Figure 4.**
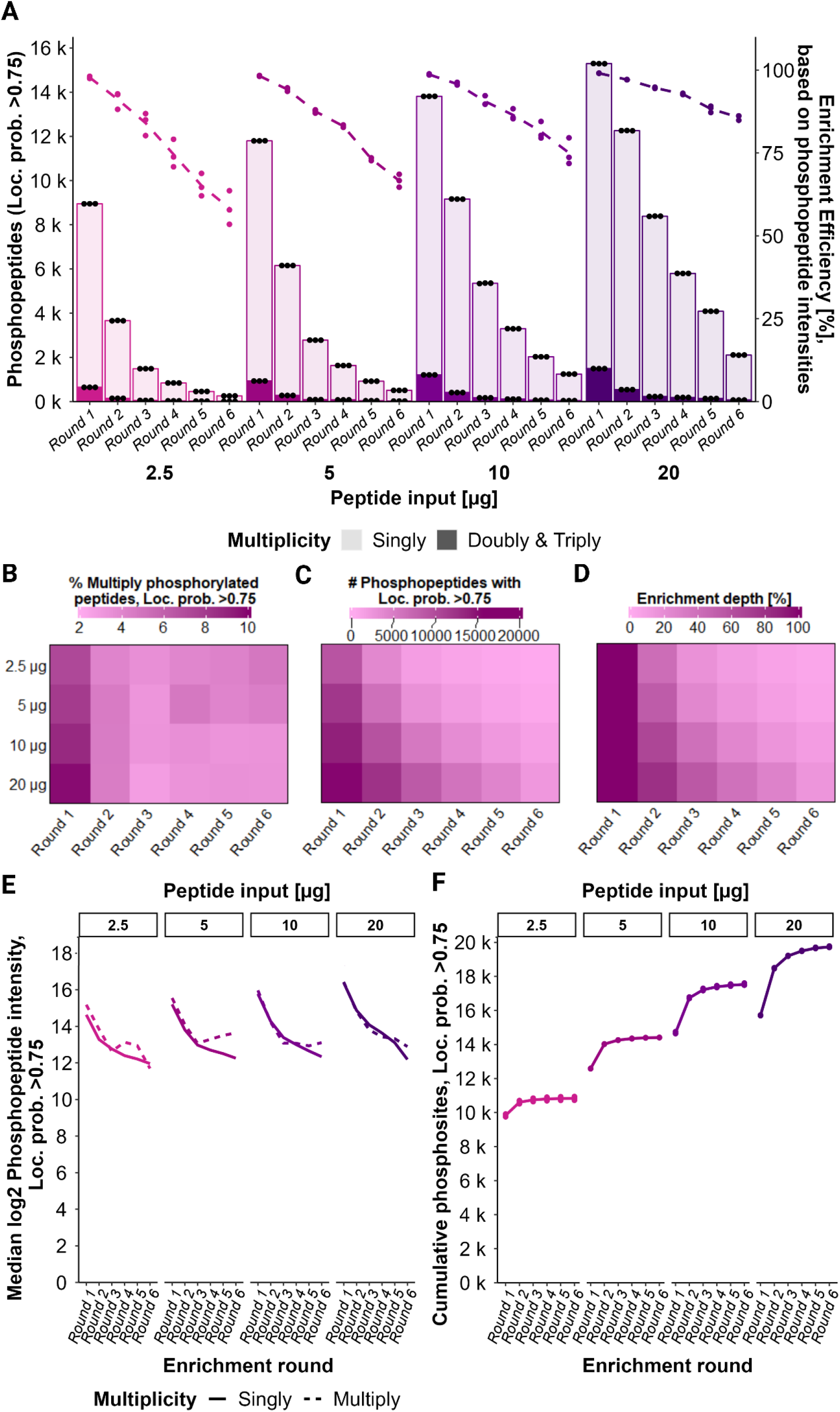
Evaluation of an extensive 6-round sequential enrichment approach. **(A)** Barplots show the mean numbers of singly phosphorylated (light color) and multiply-phosphorylated (dark color) peptides with loc. prob. >0.75 (dark color) identified across three experimental replicates using different peptide input amounts for a sequential six round enrichment. Each fraction (round) was obtained as eluate after the respective enrichment round and analyzed separately via LC-MS/MS. Each dot represents one experimental replicate. Line plots represent the mean enrichment efficiency across three experimental replicates in each round per peptide input amount based on phosphopeptide intensities in percentage. Each dot represents one experimental replicate. **(B-D)** Heatmaps show, for each peptide input amount and enrichment round, the **(B)** percentage of multiply-phosphorylated peptides with loc. prob. >0.75 in context of the total number of identified phosphorylated peptides with loc. prob. >0.75 **(C)** number of identified phosphopeptides with loc. prob. >0.75 in context of total identified phosphorylated peptides **(D)** enrichment depth in percentage in terms of number of identified phosphopeptides with loc. prob. >0.75 relative to round 1 of the respective peptide input amount. **(E)** Line plots show the medians of log2 mean intensities across replicates of singly and multiply-phosphorylated peptides with loc. prob. >0.75 identified in each enrichment round upon different peptide input amounts. **(F)** Line plots represent the mean number of cumulative phosphosites with loc. prob. >0.75 per peptide input amount and enrichment round across three experimental replicates. “Cumulative” refers to the addition of phosphosites which were not identified in the previous enrichment round(s). Each dot represents one experimental replicate.

Throughout the course of this project, we observed that the way DIA proteomics data, and in our case phosphoproteomics data in particular, is analyzed by the search engine (i.e. Spectronaut) had an impact on the identifications. In spectral library-free mode (direct-DIA) searches in Spectronaut, when several files are searched together, the information from all of them is used during the search, allowing data for spectral library inference to be obtained from one file and used during peptide identification in the other files. This effect is of special relevance when searching together high load samples and low load samples, and it is a strategy often employed in the field of single cell proteomics to boost identifications. In our experiments, searching all six fractions together using direct-DIA lead to an increase of the number of identified (phospho)peptides in all fractions (**Supplementary Figure S2E**). In contrast, when the search of each enriched fraction was done separately using the evaluation mode, most of the identified peptides are found uniquely in the first enrichment (**Supplementary Figure S2E**).

We previously observed that the site localization score decreased with lower peptide input and hence phosphopeptide intensity (**Figure 3B**). Since we observed a constant decrease in median phosphopeptide intensity with each subsequent enrichment (**Figure 4E**), we evaluated whether the site localization scores also worsened in each subsequent enrichment (**Supplementary Figure S3A-B-C-D**). Interestingly, such a trend was not observed for lower input amounts (**Supplementary Figure S3C-D**). Potentially, this could be due to the lower number of phosphopeptides identified in the last enrichment rounds when starting with 2.5 or 5 µg, which likely represents the more abundant phosphopeptides and is therefore easier to localize (**Supplementary Figure S4A**). Conversely, for higher input amounts, the population of phosphopeptides in the last enrichment might include less abundant phosphopeptides that result in worse localization scores.

Finally, we hypothesized that sequential enrichment might eventually deplete the most abundant phosphopeptides, allowing other phosphopeptide species to be enriched. Interestingly, we observed that while the overall intensity in the population of singly phosphorylated peptides decreased over time, the multiply-phosphorylated counterpart increased towards the last fractions, especially for low input amounts (**Figure 4E and Supplementary Figure S4**). Although the number of multiply-phosphorylated peptides decreased with each enrichment (**Figure 4B**), this increment in the intensity of multiply-phosphorylated peptides could be due to higher affinity of those peptides when the overall population of phosphopeptides is depleted.

Overall, the highest increase on phosphoproteome depth when doing sequential enrichment originated mostly from the second enrichment (**Figure 4D, Figure 4F**). However, doing sequential enrichment involves not only more sample preparation time, but also an increase in subsequent MS measurement time. Moreover, there is no standardized approach on how to handle multiple enrichments from a quantitative phosphoproteomics perspective, considering the high redundancy of phosphopeptides identified across the sequentially enriched fractions and that their intensity is relative towards their environment. Therefore, we hypothesized that a potential solution to benefit from the increase in depth of sequential enrichment could be achieved by pooling the fractions prior to MS analysis. We explored this possibility for highly sensitive analysis, using 2.5 and 5 µg of peptide as starting amounts for enrichment. Interestingly, we observed a gain of 7% (from 9,750 phosphopeptides in one single enrichment to 10,511 phosphopeptides in a pooled sample) when combining first and second enrichment from 5 µg of input peptides (**Figure 5A**). The gain in IDs in the pooled sample could be potentially due to a boost in the intensity (**Figure 5B**). However, we did not observe such a significant win when using 2.5 µg of peptide (only 2% increase in phosphopeptides) (**Figure 5A-B**). Furthermore, we explored the impact of pooling from a quantitative perspective by calculating the CVs of the pooled fractions and comparing it to separate enrichments or the cumulative strategy (i.e. 1st and 2nd enrichment analyzed separately by LC-MS/MS, and the resulting data merged in-silico afterwards). Reassuringly, CVs were not significantly affected by pooling the samples and remained with a median <20% (**Figure 5C**).

**Figure 5.**
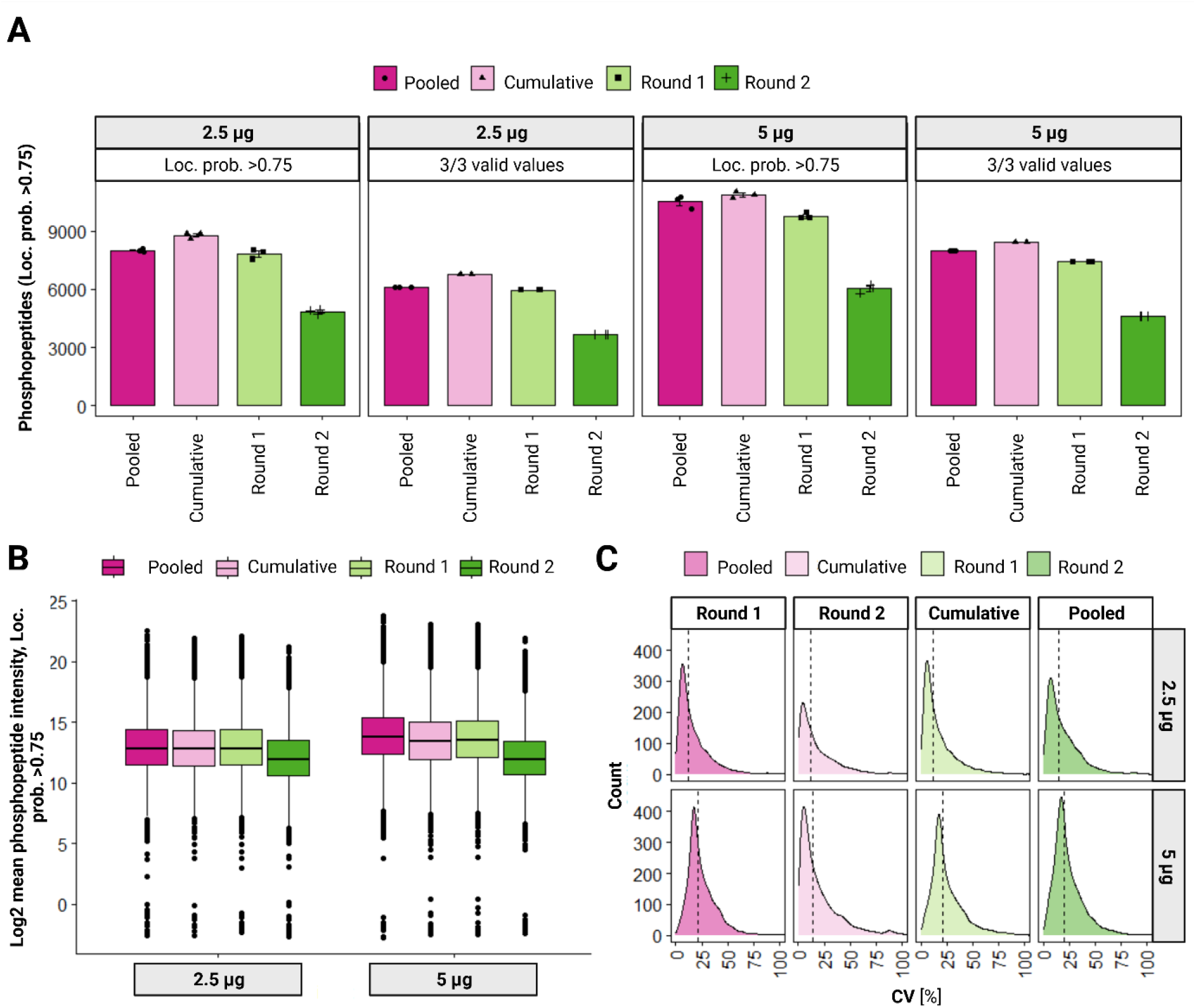
Strategies for pooling sequential enrichment samples. **(B)** Barplots show the numbers of phosphopeptides with loc. prob. >0.75 (dark color), 3/3 valid intensity values among replicates (medium color) or with a CV <0.2 among replicate intensities (light color) identified in a two round sequential enrichment approach using 2.5 µg (pink) or 5 µg (purple) peptide input amount. Each dot represents one replicate. Each fraction (round) was either obtained as eluate after the respective enrichment round and analyzed separately via LC-MS/MS (“Round 1” and “Round 2”) or obtained as a pooled eluate by reusing the elution buffer from the first enrichment round (“Pooled”). “Cumulative” refers to cumulation of unique phosphopeptide IDs identified in the separate fractions (“Round 1” & “Round 2”) during data analysis. **(C)** Boxplots show the log2 mean intensities of identified phosphopeptides with loc. prob. >0.75 per fraction and peptide input amount. **(D)** Density plots show the distribution of CVs across replicates of phosphopeptide (loc. prob. >0.75) intensities per fraction (columns) and peptide input (rows) after normalization. The labels within the density plots show the median CVs.

Next, we tried to exploit further the benefit of pooling sequential enrichment fractions by modifying the conditions of each enrichment to favor complementary populations of phosphopeptides. In particular we applied our observation that the amount of beads used inversely correlates with the number of multiply-phosphorylated peptides identified in a sample. We hypothesized that, when the amount of beads is limited, the competitive binding conditions lead to favored binding of multiply-phosphorylated peptides. Consequently, to take advantage of this in a sequential enrichment strategy, we designed the following experiment: 1st enrichment with 1 µL of beads, 2nd enrichment adding 1 µL of new beads, and 3rd enrichment adding 2 µL of new beads. We either analyzed each enrichment round fraction separately and cumulated the phosphopeptides during data analysis (cumulative approach) or pooled them into one fraction by reusing the same elution buffer aliquot (pooled approach). Additionally, we performed the standard enrichment strategy with 5 µL of beads as comparison (**Figure 6**). The results revealed that there was a significant gain when using this sequential strategy. The pooled sequential approach improved the phosphoproteome depth compared to a single enrichment in standard conditions when starting with input amounts of at least 15 µg (from 9,247 to 11,356 phosphopeptides for 15 µg, and from 12,568 to 14,085 phosphopeptides for 30 µg). Moreover, the pooled approach yielded more phosphopeptide IDs than the cumulative approach for all input amounts (6,385 vs. 5,343 phosphopeptides for 5 µg, 11,356 vs. 10,442 phosphopeptides for 15 µg and 14,085 vs. 13,004 phosphopeptides for 30 µg). Interestingly, we observed that there was no such gain when using lower input amounts (i.e. 5 µg). This might indicate that the beads-to-peptide ratio was not optimized for such low amounts, and that more optimization might be required to make this strategy beneficial. Overall, we were able to confirm that sequential enrichment approaches can significantly increase phosphopeptide identifications compared to a standard one-round enrichment and observed that pooling fractions from multiple sequential enrichment rounds can outperform their separate analysis in terms of phosphopeptide identifications.

**Figure 6.**
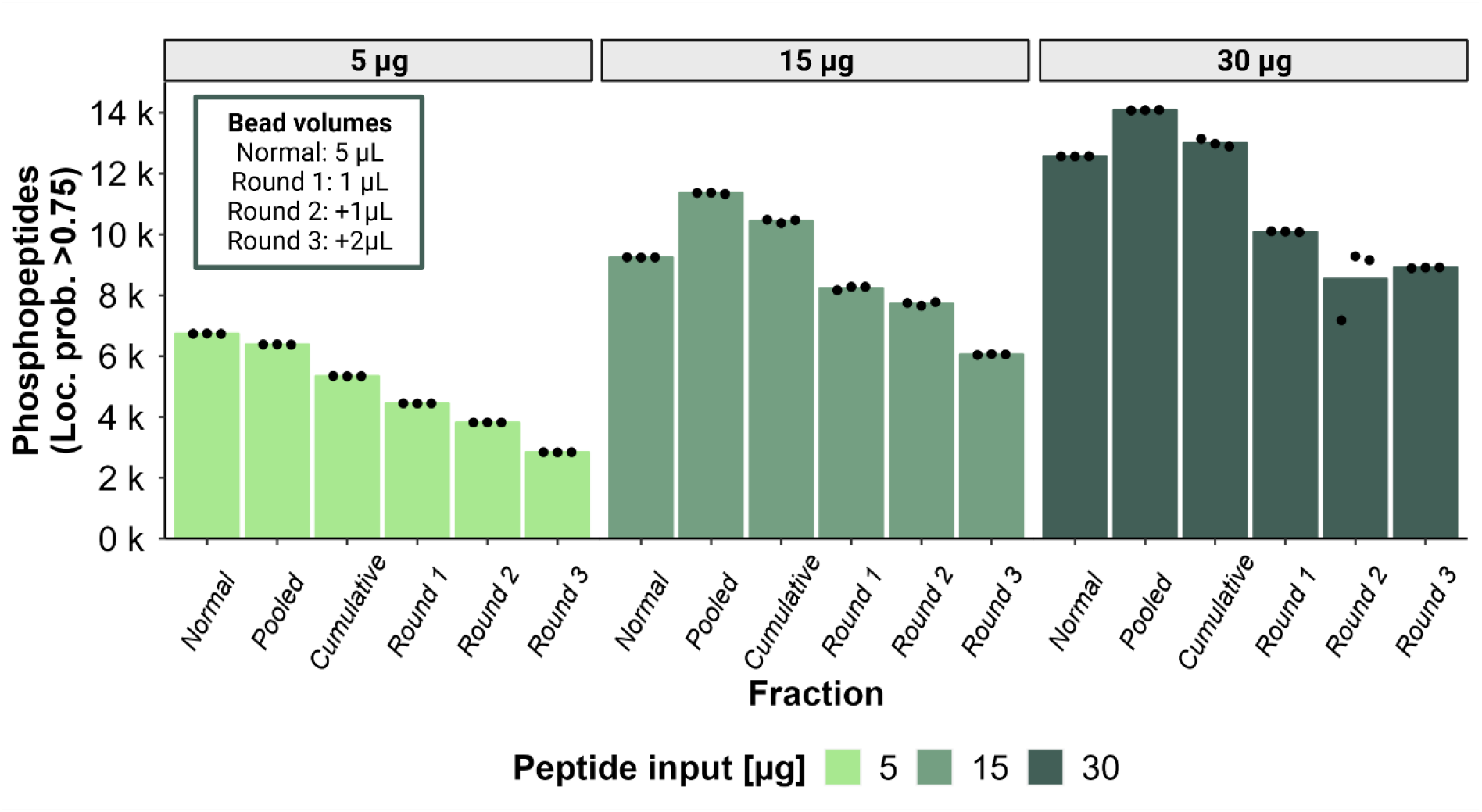
Refined sequential enrichment pooling approach with increasing Zr-IMAC HP bead volume. Barplots show the numbers of phosphopeptides with loc. prob. >0.75 identified using a three-round sequential enrichment approach with increasing Zr-IMAC HP bead volume for different peptide input amounts. “Normal” represents a standard single-round enrichment with 5 µL beads. “Pooled” represents a sequential enrichment for three rounds with increasing bead volume (Round 1: 1 µL beads, Round 2: +1 µL beads, Round 3: + 2 µL beads) and rounds pooled into the same elution buffer. “Round 1”, “Round 2” and “Round 3” represent the identifications in the respective separately collected and analyzed fractions. “Cumulative” refers to cumulation of unique phosphopeptides identified in the separate fractions (“Round 1”, “Round 2”, “Round 3”) during data analysis.

### Combination of sequential enrichment with LC-MS/MS analysis on an Orbitrap Astral Mass Spectrometer

Finally, the evolution of mass spectrometers is one of the most significant aspects of phosphoproteomics leading to improvements in sensitivity, speed and depth of analysis. Therefore, we decided to complete our systematic evaluation by benchmarking the optimized phosphoproteomics workflow using the latest-generation high-end proteomics-grade MS instrumentation, the Orbitrap Astral mass spectrometer.

To evaluate the performance of the Orbitrap Astral for phosphoproteomics using the optimized phosphopeptide enrichment workflow in settings best representing typical biological experiments, we performed the phosphopeptide-enrichment starting from different numbers of HeLa cells. Consequently, the resulting phosphoproteome coverage reflects how sensitive the protocol is to the number of input cells. Our experiment used four replicates for six different number of cells, ranging from 1 million cells to 10,000 cells. The different cell samples were lysed in SDS-buffer and digested using PAC-based trypsin digestion, followed by phosphopeptide enrichment without any desalting step to minimize losses. The phosphopeptide enrichment was performed using the optimized parameters described in this project. Two rounds of sequential enrichment were performed, eluates were pooled together for analysis and EvoTipped samples were analysed by a 40-SPD Whisper flow method with half-an-hour LC gradient time on the Orbitrap Astral using narrow-window data-independent acquisition (nDIA) with narrow DIA isolation windows^9^.

This resulting data reflects the higher sensitivity of the Orbitrap Astral MS with more than 35,000 phosphopeptides from 1M cells or >32,000 phosphopeptides mapping to 26,000 class I phosphosites quantified in at least two samples when starting with 0.5 million cells (**Figure 7B**). This is approximately equivalent of 34 µg of purified tryptic peptides, which is an improvement of approx. two-fold when compared to the previous coverage achieved from 30µg of purified peptides in the Orbitrap Exploris 480 with up to 16,500 phosphopeptides (**Figure 2**). The Orbitrap Astral allows a comprehensive coverage of the phosphoproteome for amounts as low as 50,000 cells (7,967 phosphopeptides). With less cells, the output dropped below the sensitivity limits of our workflow (**Figure 7A-B**). The enrichment efficiency calculated as a function of overall intensities was >90% as expected (**Supplementary Figure S5**). We also evaluated the site localization scores obtained from the phosphopeptides, and observed a similar trend as the one observed for Orbitrap Exploris 480 data (**Figure 3B**) with a clear drop in localization scores for lower cell inputs (**Figure 7C**).

**Figure 7.**
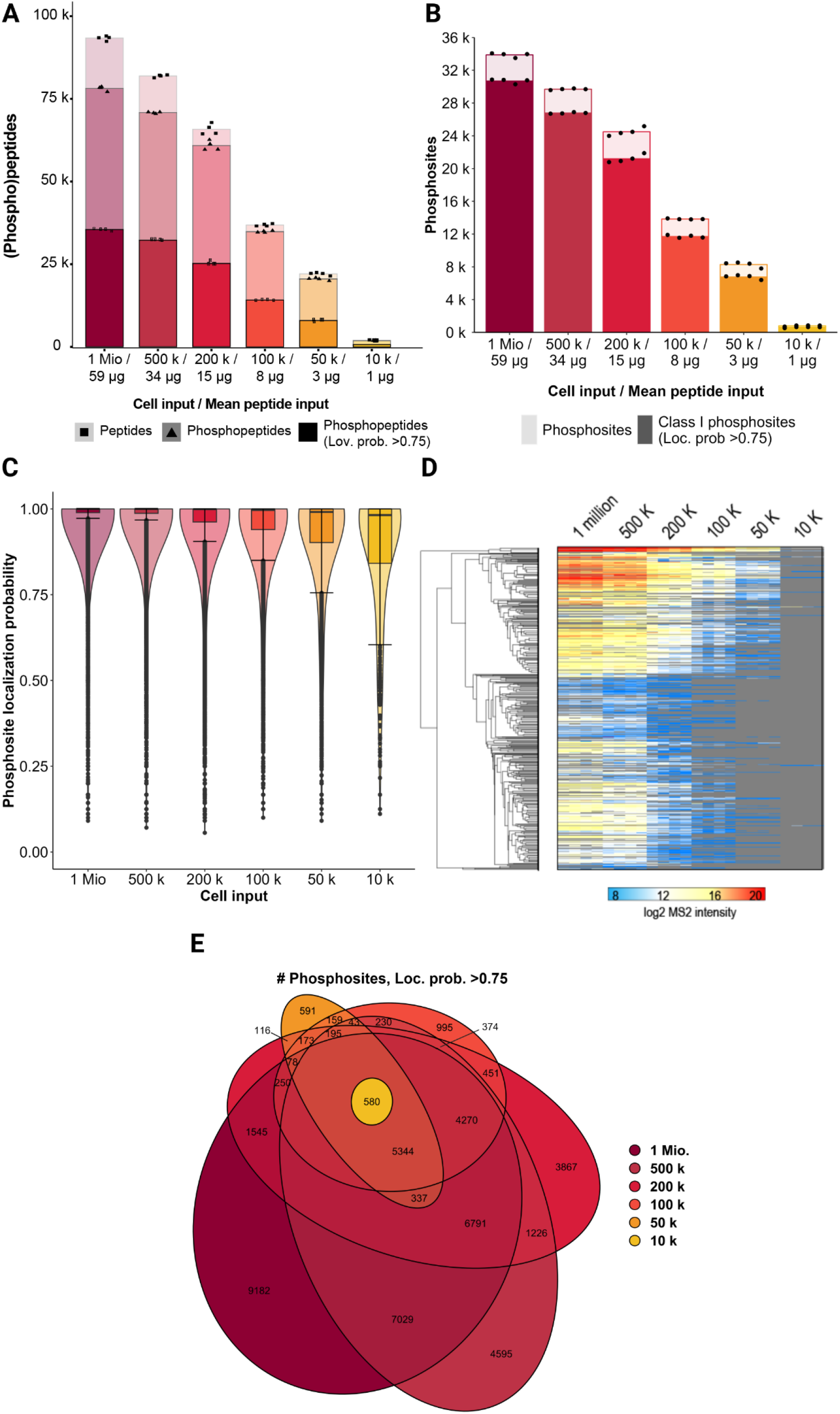
Employment of a 2-round sequential enrichment pooling approach for analysis of a HeLa dilution series in combination with LC-MS/MS analysis on an Orbitrap Astral Mass Spectrometer. **(A)** Barplots show the mean numbers of peptides (light color), phosphopeptides (medium color) and phosphopeptides with loc. prob. >0.75 (dark color) identified across four experimental replicates using different cell input amounts in a 2-round pooled sequential enrichment. Each dot represents one experimental replicate. **(B)** Barplots show the mean numbers of phosphosites (light color), and class I phosphosites (loc. prob. >0.75) (dark color) identified across four experimental replicates using different cell input amounts in a 2-round pooled sequential enrichment. Each dot represents one experimental replicate. **(C)** Violin plots show the range and distribution of the localization probability of phosphosites identified using different cell input amounts. **(D)** Heatmap shows the log2 mean intensities of unique phosphopeptides identified across four experimental replicates using different cell input amounts in a 2-round pooled sequential enrichment. **(E)** The venn diagram shows uniquely and commonly identified phosphosites with loc. prob. >0.75 among different cell input amounts in a 2-round pooled sequential enrichment.

## Conclusions

Our extensive optimization of phosphopeptide enrichment conditions elucidated that the key parameters, including beads-to-peptide ratio, binding time, competitive-binder concentration and sample volume, markedly influence phosphopeptide enrichment efficiency, phosphopeptide recovery and phosphosite localization scores. Particularly, our findings underscore that multiply-phosphorylated peptides exhibit enhanced affinity towards Zr-IMAC HP beads, leading to their preferential enrichment under competitive binding conditions such as high glycolic acid concentration or low bead volumes. We recommend to adapt enrichment conditions accordingly to the specific needs when aiming for either highest phosphoproteome depth, enrichment efficiency, or proportion of multiply-phosphorylated peptides.

We propose sequential phosphopeptide enrichment as a powerful strategy to further amplify the depth of phosphopeptide analysis. Our study indicates that while initial enrichment rounds demonstrate higher enrichment efficiency, subsequent rounds do not, providing only minimal improvements in phosphoproteome depth after the third round. Therefore, a sequential approach with two enrichment rounds seems to be favorable for most applications in a range from 20 µg to 2.5 µg of peptide input, although more rounds might offer further improvement for high peptide input amounts.

Importantly, the post-acquisition analysis in the search engine (i.e. Spectronaut) of separate fractions or experimental conditions has so far been the standard method for method optimization. However, in the case of sequential enrichment, the data analysis of one phosphopeptide entity with multiple LFQ intensities derived from independent enrichments is not trivial. It can result in higher variability and/or increased CVs, hindering subsequent data interpretation, especially when performing label-free quantification. Our data shows that pooling fractions into a single LC-MS/MS analysis is a good alternative to circumvent these issues, while decreasing the LC-MS/MS analysis time. In this regard, we present an improved strategy based on incremental addition of beads and subsequent fraction pooling, which offers up to 20% boosted phosphoproteome coverage compared to standard enrichment, while maintaining high sample-throughput and straightforward data analysis.

Finally, we report that our optimized phosphoproteomics pipeline can be translated to the newest generation of mass spectrometers, such as the Orbitrap Astral, that can increase the phosphopeptide coverage by 2x, allowing for deep phosphoproteomics analysis without the need to scale up the starting cell amounts.

Future research might explore refinement of enrichment conditions that simultaneously maximize both enrichment efficiency and phosphoproteome depth. Combining optimal enrichment conditions with strategic application-tailored pooling of sequential enrichments and could pave the way for more comprehensive phosphoproteomics investigations.

## Acknowledgements

The authors would like to thank Justin Jordaan, Isak Gerber and Stoyan Stoychev (ReSyn Biosciences) for their valuable input on experimental design and results. Work at The Novo Nordisk Foundation Center for Protein Research (CPR) is funded in part by a donation from the Novo Nordisk Foundation (NNF14CC0001). This work has also been funded as part of EPIC-XS project under the grant agreement no. 823839 funded by the Horizon 2020 programme of the European Union. P.B. was supported by the “International Exchange Program” (Vienna Doctoral School in Chemistry, DoSChem), the funding program “Internationale Kommunikation” (Österreichische Forschungsgemeinschaft, ÖFG) and the “Erasmus+ Traineeship Mobility” program. C.K. is supported by the Marie Skłodowska Curie European Training Network “PUSHH” (grant number No. 861389).

## Author contributions

A.M.-V. and J.V.O. designed the experiments. C.K. helped in the experimental design and optimization of the phosphopeptide enrichment protocol. P.B. performed phosphoproteomics experiments. A.M.-V. and P.B. analyzed the data and generated figures. I.P. prepared, measured and analyzed the samples in the HeLa cell dilution series experiment. J.V.O. and C.G. and critically evaluated results. A.M.-V., J.V.O., P.B. and I.P. wrote the manuscript. All authors read, edited and approved the final version of the manuscript.

## Supplementary figures

**Supplementary Figure S1.**
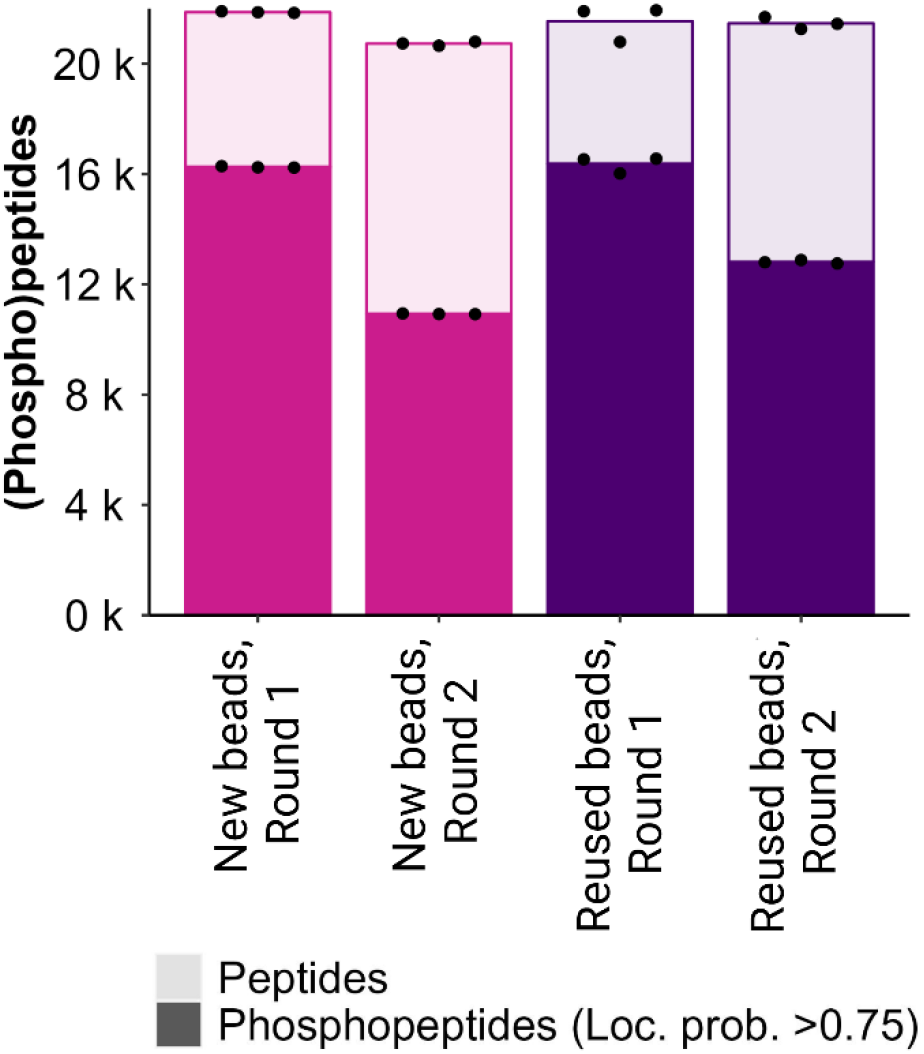
Effect of bead reuse or exchange in a sequential phosphopeptide enrichment approach. **(A)** Barplots show the mean numbers of peptides (light color) or phosphopeptides with loc. prob. >0.75 (dark color) identified across three experimental replicates in a sequential enrichment approach with two rounds in which the beads were either exchanged after the first round (pink) or reused (purple). The peptide input amount for the enrichment was 30 µg for all conditions. Each dot indicates one experimental replicate. **(B)** Barplots show the numbers of phosphopeptides with loc. prob. >0.75 (dark color), 3/3 valid intensity values among replicates (medium color) or with a CV <0.2 among replicate intensities (light color) identified in a two round sequential enrichment approach either exchanging (pink) or reusing (purple) the beads from the first enrichment round. Each dot represents one experimental replicate.

**Supplementary Figure S2.**
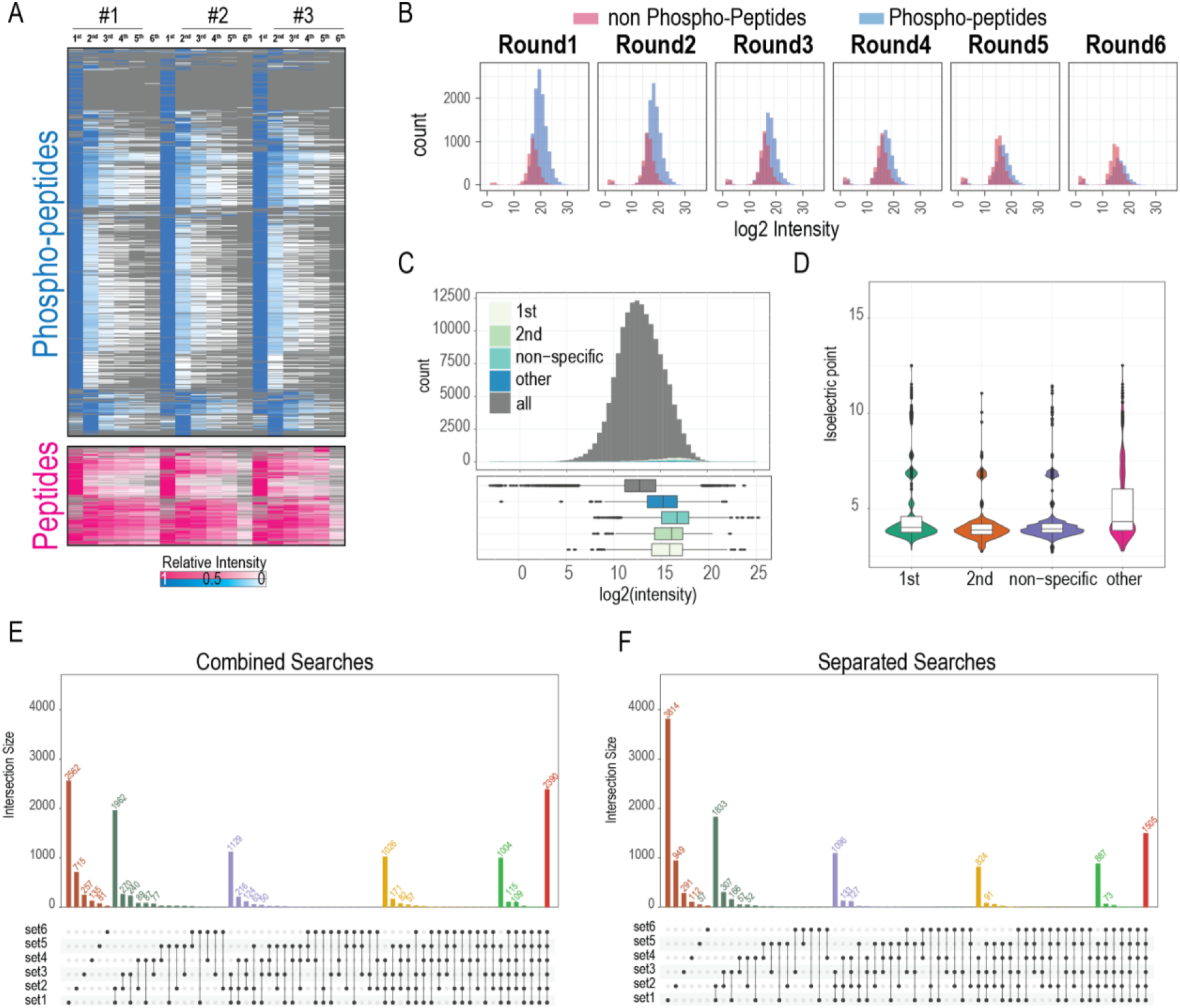
(Phospho)peptide elution profiles across fractions upon sequential enrichment with 6 rounds. **(A)** Elution profile of phosphopeptides (blue) and non-phospho peptides (pink) across six sequential enrichments using 20 µg of peptide input in three separate experimental replicates. The intensity plotted is scaled from 1 to 0 across the six sequential enrichments. **(B)** Histograms showing the peptide intensity (in log2) distribution for one experiment and six sequential enrichments. In blue: phosphopeptides, in pink: non-phospho peptides. **(C)** Histogram (top) and boxplot (bottom) showing the whole peptide intensity distribution (gray) of the whole proteome of A549 (analyzed as a single-shot in Orbitrap-Astral, data from Guzman et al^9^). highlighted in blue colors, the distribution of the non-phosphorylated peptide intensities found in the six sequential enrichment experiments when measured in a whole proteome. The different categories (1st, 2nd, non-specific and other) correspond to different elution profiles of the non-phosphopeptides as observed in panel A. 1st: peptides eluting mainly in the 1st enrichment. 2nd: peptides eluting mainly in the second enrichment. Non-specific: peptides showing a constant elution across the first enrichments. Other: other peptides with no pattern in their elution. **(D)** Isoelectric point distribution values in the non-phosphopeptides measured in the six enrichment experiments shown in panel A. Categories are the same as described in panel C. **(E-F)** Effect of search strategy on the sequential enrichment results. **(E)** Results of overlap in identifications between enrichments when the samples are analyzed together in Spectronaut. **(F)** Results of overlap in identifications between enrichments when the samples are analyzed separately in Spectronaut.

**Supplementary Figure S3.**
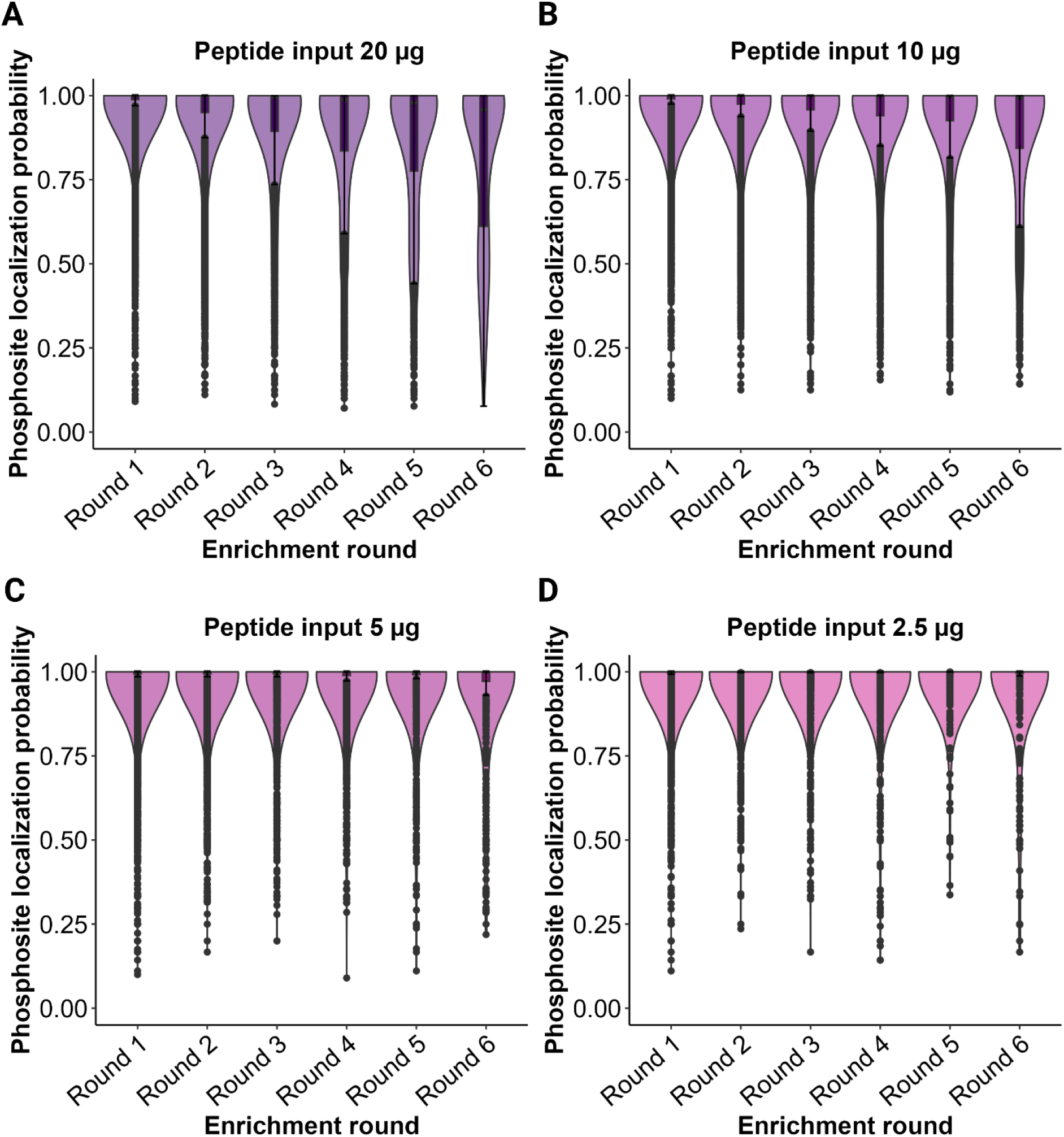
Phosphosite localization in sequential enrichment approach with 6 rounds. **(A-D)** Violin plots show the range and distribution of the localization probability of phosphosites identified in each enrichment round of a 6 round sequential enrichment, using different peptide input amounts (A: 20 µg, B: 10 µg, C: 5 µg, D: 2.5 µg)

**Supplementary Figure S4.**
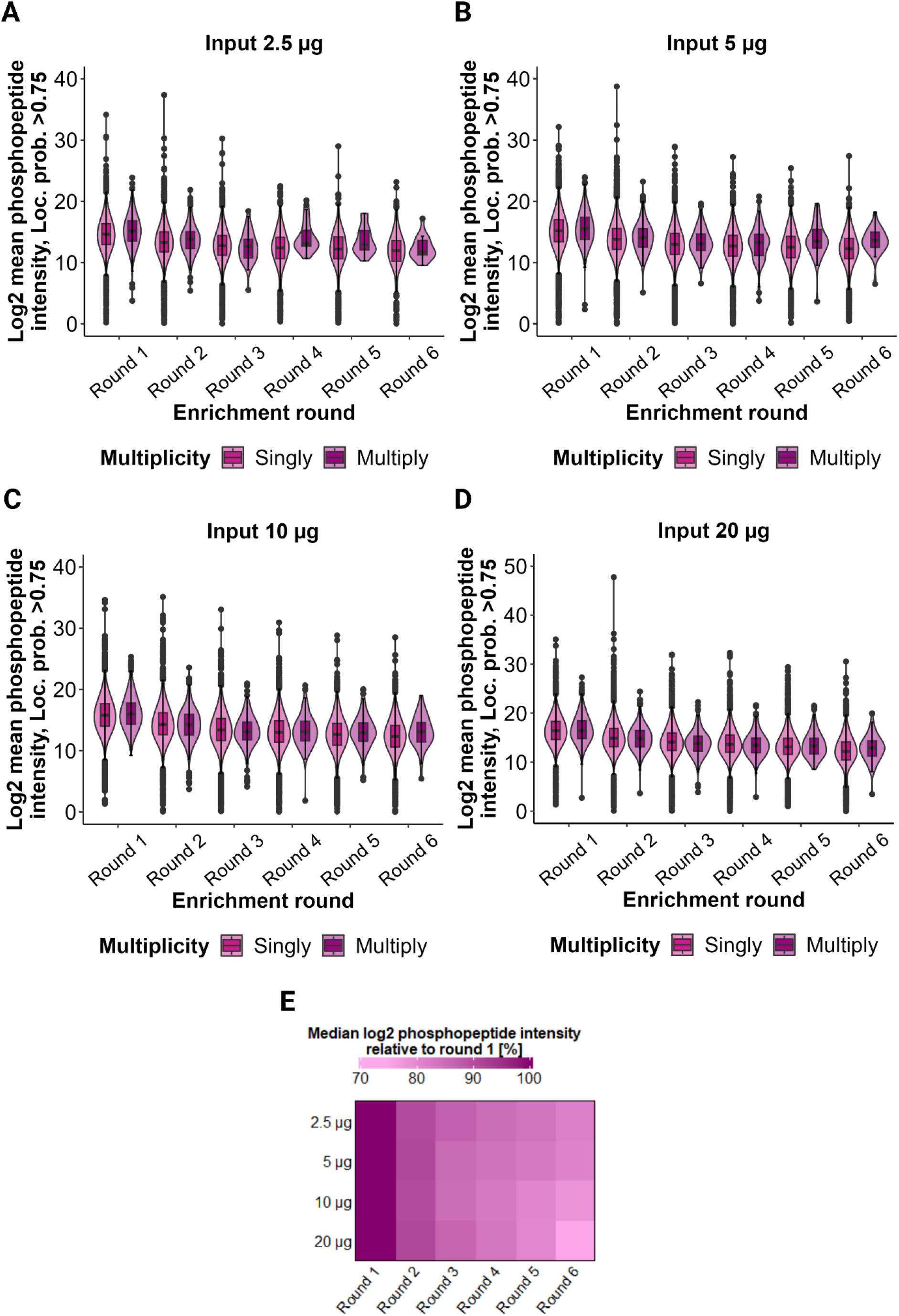
Phosphopeptide IDs in a sequential enrichment approach with 3 rounds and different bead-binding times. **(A-D)** Violin plots show the distribution log2 mean phosphopeptide intensities of singly and multiply phosphorylated peptides in each enrichment round of a 6 round sequential enrichment, using different peptide input amounts (A: 20 µg, B: 10 µg, C: 5 µg, D: 2.5 µg). (**E**) Heatmap shows the median log 2 phosphopeptide intensities of singly and multiply phosphorylated peptides relative to round 1 of a 6 round sequential enrichment, using different peptide input amounts.

**Supplementary Figure S5.**
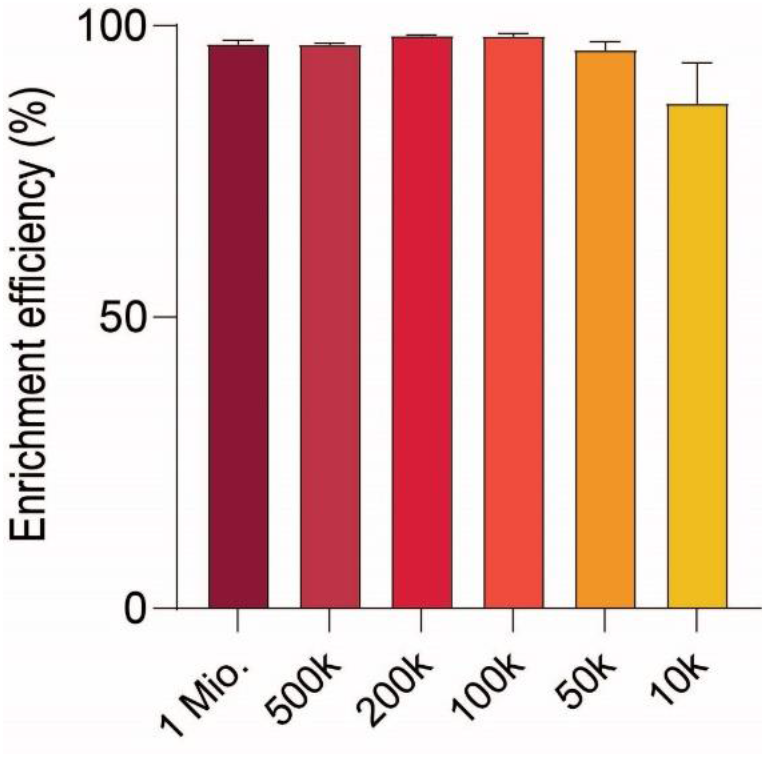
Phosphopeptide enrichment efficiency calculated from peptide intensity. Barplots show the average of the phosphopeptide enrichment efficiency of data obtained using the Orbitrap Astral with different cell amounts as input in the workflow.

